# Rapid chromosomal rearrangements and sex chromosome turnover underlie the evolution of parapatric Pacific *Scomber* mackerels

**DOI:** 10.64898/2026.04.11.717877

**Authors:** Ahammad Kabir, Ryosuke Yazawa, Dana Marielba Silva, Jorge M.O. Fernandes, Masaomi Hamasaki, Sota Yoshikawa, Hiroaki Suetake, Kiyoshi Kikuchi, Sho Hosoya

## Abstract

Teleost display remarkable species diversity despite relatively conserved karyotypes, suggesting an important role for chromosomal rearrangements in speciation. Yet this hypothesis remains poorly tested in pelagic marine fishes, as genomic studies have predominantly focused on freshwater and coastal taxa. In this study, we investigated the genomic basis of hybrid incompatibility between two parapatric Pacific mackerels, *Scomber japonicus* and *S. australasicus*, hypothesizing chromosomal rearrangements as the major driving forces. We generated haplotype-resolved de novo genome assemblies for both Pacific species and assessed genomic rearrangements along with the genome of the Atlantic species, *S. scombrus,* reconstructed from publicly available data. Comparative genomic analyses revealed extensive chromosomal rearrangements across the genomes. Notably, the rate of chromosomal inversions was approximately sevenfold higher between the two Pacific species than between allopatric lineages. These rearrangements included complex structural changes involving megabase-scale inversions and associated translocations. Our analyses also showed that the sex chromosomes of the three species evolved independently. In the two Pacific species, large recombination-suppression regions (>10 Mb) arose convergently but through distinct mechanisms: tandem chromosomal inversions in *S. japonicus* and, most likely, transposable element–mediated sequence divergence in *S. australasicus*. By contrast, recombination suppression in *S. scombrus* is restricted to a ∼14 kb hemizygous region containing *amhr2Y*, generated by duplication and translocation. Population-level analyses further revealed ongoing evolution of recombination-suppressed regions in the Pacific species and uncovered multiple Y chromosome lineages in *S. australasicus* that differ markedly in the extent of recombination suppression. Together, these results demonstrate rapid structural genome evolution in *Scomber* and provide a genomic framework for understanding how chromosomal rearrangements contribute to reproductive isolation, sex chromosome turnover, and the diversification of pelagic marine fishes.

## Introduction

Teleost fishes exhibit extraordinary species richness and have diversified into a broad range of ecological niches^1–3^. Nevertheless, they present relatively conserved karyotypes, with limited variation in chromosome number across species^4^. This discrepancy suggests that chromosomal rearrangements associated with large-scale structural variations (SVs) may have played important roles in speciation and the maintenance of species boundaries. Advanced short-read sequencing technology has highlighted the contributions of variations in gene sequences, i.e., single nucleotide polymorphisms (SNPs), to species divergence and local adaptation^5–7^. Long-read and long-range sequencing technologies are now used to elucidate the importance of large-scale SVs, such as gene duplications, deletions, inversions, and translocations, in shaping reproductive barriers and ecological differentiation^8–10^.

Scombridae, a group of epipelagic migratory fish that includes mackerels and tuna, are recognized as ecologically and economically important because of their global distribution and worldwide consumption. Notably, four of the top ten marine fish species in global capture fisheries production belong to this family (Table 7 of The State of World Fisheries and Aquaculture, FAO, 2024). Many Scombridae species share a conserved karyotype (2n = 48, predominantly acrocentric chromosomes)^11,12^, yet the genus *Scomber* has undergone relatively recent and rapid diversification into at least four closely related species distributed across the Atlantic, Pacific, and Indian Oceans. Among these species, the two Pacific species, chub mackerel (*S. japonicus*) and blue mackerel (*S. australasicus*), which exhibit parapatric distributions around Japan and the South China Sea, have diverged within approximately two million years (Supplementary Fig. 1). As their spawning area and timing overlap^13^, natural hybrids occur between these species^14,15^; however, they are mostly infertile^16^. This finding suggests that hybrid incompatibility has evolved between the two species within a relatively short time, even though interspecific hybridization is known to occur in other fish lineages despite much deeper divergence times^17–19^. Intriguingly, the two Pacific *Scomber* species also possess distinct sex determination systems: female heterogamety (ZW) in *S. japonicus* and male heterogamety (XY) in *S. australasicus*^20^, suggesting that the evolution of their sex chromosomes contributes to reproductive isolation. In parallel, large-scale SVs may also play a role in species divergence and the maintenance of species boundaries. Comparisons of the genomes of these two rapidly speciated species and clarification of chromosome rearrangements at the whole-genome level, including sex chromosomes, are expected to provide important insights into genome evolution, speciation, and species maintenance.

In this study, we present high-quality haplotype-resolved *de novo* genome assemblies for *S. japonicus* and *S. australasicus* constructed using PacBio Revio long-read sequencing and Dovetail Omni-C long-range data. We further perform low-coverage whole-genome resequencing (lcWGR) across multiple populations of both species to investigate SVs at the population scale. Public genomic data from the Atlantic species *S. scombrus*, including long-read and long-range information, as well as RAD-seq data obtained from wild populations, are also incorporated as an outgroup. By identifying chromosomal inversions and their flanking repeat structures, we explore the mechanisms driving genomic rearrangement that may be associated with hybrid incompatibility. Additionally, we identify and characterize previously unknown sex chromosomes to better understand their roles in speciation and species maintenance. Our findings provide insights into the genomic basis of species maintenance in parapatric species groups and highlight the potential of structural genomic resources for the sustainable management of pelagic fisheries, which are becoming increasingly vulnerable to long-term environmental fluctuations^21–23^.

## Results

### Haplotype-resolved reference genomes of three *Scomber* species

We generated haplotype-resolved genome assemblies for ZW *S. japonicus* (DDBJ BioProject Accession: **********) and XY *S. australasicus* (**********) (Table 1, Supplementary Figs. 2 and 3). More than 98% of the assembled sequences were anchored into 24 superscaffolds per haploid genome, consistent with the known karyotype, yielding assemblies of 823.7–838.7 Mb (scaffold N50 = 35 Mb) with BUSCO completeness of > 97% (Actinopterygii). As the current reference genome of *S. japonicus* (fScoJap1.pri) was generated from a ZZ individual, our assembly represents the first ZW genome for this species and achieves a comparable level of completeness. We located telomeric regions characterized by an array of vertebrate telomeric repeats (TTAGGG) ^24^ in at least one of the two haploid genomes for 20 superscaffolds of the *S. japonicus* genome and 17 superscaffolds of the *S. australasicus* genome (Supplementary Figs. 2B and 3B, Supplementary Table S1). These results underscore the (nearly) telomere-to-telomere (T2T) quality of the *S. japonicus* and *S. australasicus* genome assemblies. Within these genomes, we identified approximately 29,000 and 24,000 protein-coding genes for *S. japonicus* and *S. australasicus*, respectively (Table 1, Supplementary Table S2). The genomes contained 27 to 28% repetitive sequences; 20% were transposable elements (TEs), and the rest consisted of simple repeats, satellite repeats, and low-complexity regions.

**Table 1.** Assembly statistics for the haplotype-resolved genomes of a female *S. japonicus* (SJF), a male *S. australasicus* (SAM) and a male *S. scombrus* (SSM)

|  | *S. japonicus* |  | *S. australasicus* |  | *S. scombrus* |  |
| --- | --- | --- | --- | --- | --- | --- |
| Sex | female (ZW) |  | male (XY) |  | male (XY*) |  |
| Long reads (Gb) | REVIO HiFi (76.82) |  | REVIO HiFi (51.13) |  | Sequel II HiFi (19.92) |  |
| 3C-methods (Gb) | Omni-C (121.31) |  | Omni-C (137.58) |  | Arima Hi-C (106.96) |  |
| Haploid genome name | SJF-hap1 | SJF-hap2 | SAM-hap1 | SAM-hap2 | SSM-hap1 | SSM-hap2 |
| Assemble statistics |  |  |  |  |  |  |
| Number of scaffolds | 28 | 35 | 41 | 127 | 569 | 435 |
| Largest (bp) | 43,470,709 | 44,519,000 | 42,926,962 | 42,750,987 | 38,912,451 | 38,599,483 |
| Shortest (bp) | 250,000 | 5,000 | 3,000 | 17,696 | 4,366 | 6,274 |
| Average (bp) | 29,746,910 | 23,963,238 | 20,226,707 | 6,486,018 | 1,339,128 | 1,715,061 |
| N50 (bp) | 35,452,366 | 35,330,575 | 35,519,200 | 34,591,313 | 31,183,350 | 31,631,483 |
| Number of superscaffolds (chromosome) | 24 | 24 | 24 | 24 | 24 | 24 |
| Genome size (bp) | 832,913,979 | 838,714,842 | 829,293,995 | 823,725,848 | 761,937,010 | 754,896,464 |
| Chromosomes with telomeres (both end) | 13 | 11 | 12 | 10 | 0 | 0 |
| Chromosomes with telomeres (one end) | 10 | 12 | 8 | 11 | 8 | 11 |
| Chromosomes without telomeres | 1 | 1 | 4 | 3 | 16 | 13 |

| BUSCO score |  |  |  |  |  |  |
| --- | --- | --- | --- | --- | --- | --- |
| Single copy | 97.2% | 97.3% | 97.1% | 97.3% | 96.5% | 96.7% |
| Duplicated | 1.0% | 0.9% | 1.0% | 0.9% | 1.2% | 1.0% |
| Fragmented | 0.6% | 0.7% | 0.6% | 0.5% | 0.7% | 0.7% |
| Missing | 1.2% | 1.1% | 1.3% | 1.3% | 1.6% | 1.6% |
| Protein-coding genes | 28,913 | 28,393 | 24,842 | 23,923 | 22,929 | 22,929 |
| GC contents | 40.09% | 40.05% | 40.16% | 40.12% | 39.80% | 39.75 |
| Fraction of repeat-rich regions | 28.21% | 28.19% | 26.98% | 26.78% | 25.75% | 25.97% |
| Fraction of TEs | 19.79% | 19.85% | 19.59% | 19.57% | 20.11% | 20.35% |

\*The sex of the SSM sample was inferred from the genetic analysis in this study.

The haploid-resolved genome of *S. scombrus* was reassembled using publicly available data, including PacBio HiFi long reads and Arima Hi-C data, obtained from an XY individual (BioSample: SAMEA110450232) (Supplementary Fig. 4). Although a genome assembly for this individual is already available (fScoSco1.1; GCF_963691925.1), we reassembled it using the same pipeline applied to the two Pacific species to enable direct genome comparisons without confounding effects of the assembly methodology. The total assembly sizes of the 24 major superscaffolds were 704.9–709.4 Mb (scaffold N50 = 35 Mb) per haploid genome with BUSCO completeness of > 97%. For these genomes, we could locate telomeric repeats at one end of 8 superscaffolds in the primary haplotype and 11 superscaffolds in the alternate haplotype. The overall assembly quality of the resulting genomes was comparable to that of the NCBI reference genome for this species. For gene annotation, we adopted the annotation file for fScoSco1.1 and added it to the reassembled genome. Comparable values were observed for the number of protein-coding genes and other genomic features in this species relative to those in the other two species. The distributions of TEs and simple/satellite repeats in the genomes of the three species are presented in Supplementary Fig. 5.

Hereafter, the primary and secondary haploid assemblies of *S. japonicus*, *S. australasicus*, and *S. scombrus* are referred to as hap1 and hap2, with prefixes indicating species and phenotypic sex: SJF-hap1, SJF-hap2, SAM-hap1, SAM-hap2, SSM-hap1, and SSM-hap2. Superscaffolds are referred to as chromosomes.

### Centromere positioning and karyotype inference

Centromere position is a key determinant of chromosome morphology, and its identification is essential for understanding the evolution of chromosome structure. By combining analyses of the centromeric repeat content and DNA hypomethylation patterns^25,26^, we identified the genomic locations of centromeres on at least one of the two haploid assemblies for all 24 chromosomes in *S. japonicus* and *S. australasicus* (Supplementary Table S1). In *S. japonicus*, centromeres were identified on all 24 chromosomes in both haploid genomes (Supplementary Fig. 6). In *S. australasicus*, centromeres were identified on 21 chromosomes (excluding Chr4, Chr11, and Chr24) in the primary assembly (SAM-hap1) and on 23 chromosomes (excluding Chr12) in the alternative assembly (SAM-hap2) (Supplementary Fig. 7). These centromeric and hypermethylated regions share seven conserved satellite repeat motifs (Supplementary Table S3), supporting the accuracy of centromere identification, as centromeres often contain conserved satellite DNA^27^. An analysis of chromosome arm length ratios (long arm to short arm) indicated that both species possess only mono-armed chromosomes; *S. japonicus* has 3 subtelocentric and 21 acrocentric chromosomes, whereas *S. australasicus* has 2 subtelocentric and 14 acrocentric and 8 acrocentric/telocentric chromosomes (Supplementary Table S1), largely consistent with the findings of previous cytogenetic studies^12,28^.

In contrast to the two Pacific *Scomber* species, centromeric regions could not be identified in the *S. scombrus* genome. The relatively low coverage of long HiFi reads in this species limited the contiguity of the assembly, impeding the reconstruction of complex genomic regions. Nevertheless, a synteny analysis among the three species suggested that the chromosomes of *S. scombrus* are likely acrocentric or telocentric (Fig. 1B).

**Fig. 1:**
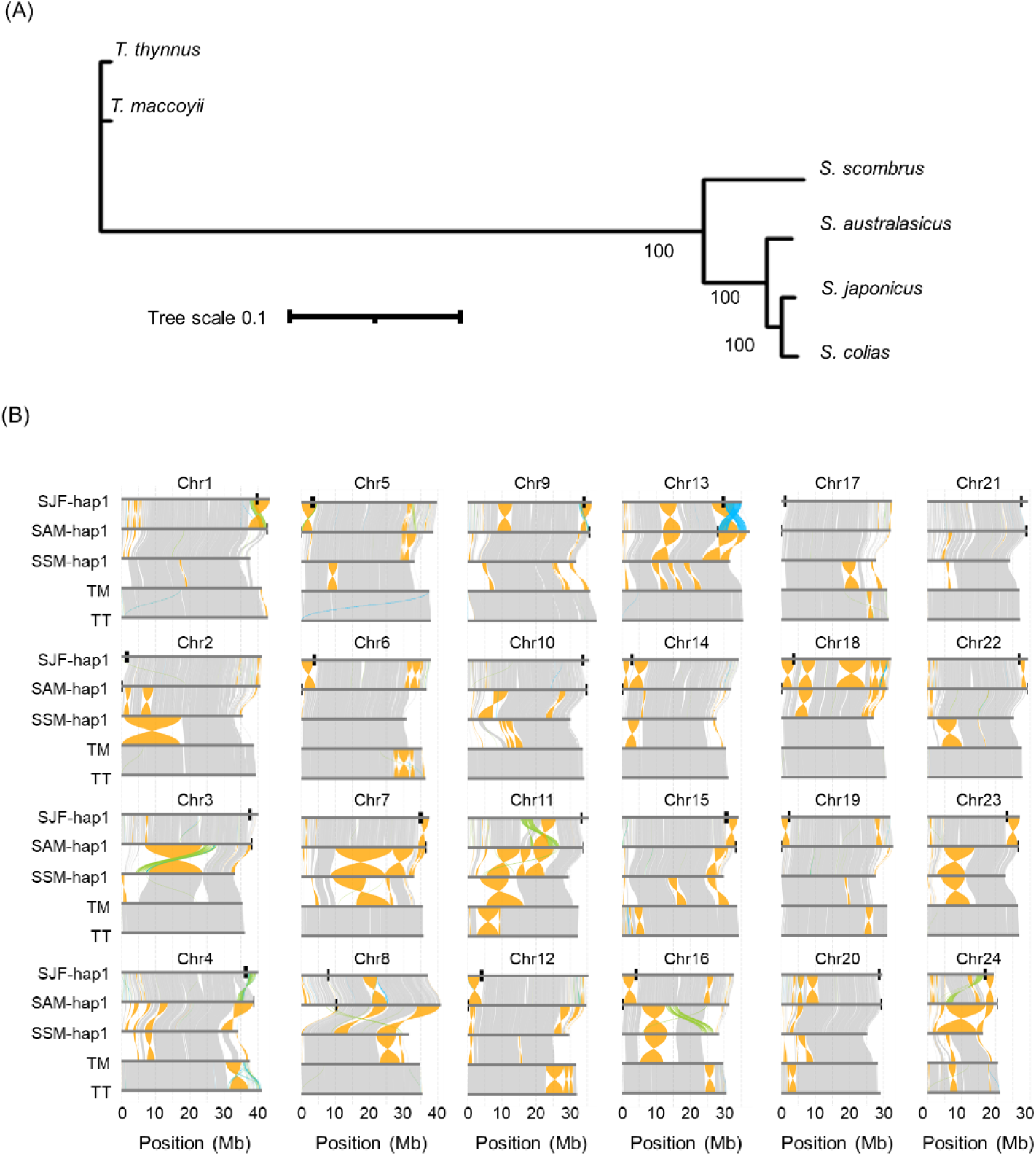
Species tree and chromosomal rearrangements among *Scomber* species and two outgroup species. **(A)** Phylogenetic relationships among two Pacific *Scomber* species and one Atlantic *Scomber* species (*S. japonicus*, *S. australasicus*, and *S. scombrus*) inferred from 6,400 single-copy orthologues together with *S. colias* and the two outgroup species *Thunnus thynnus* and *T. maccoyii*. **(B)** Comparative analyses of synteny among the primary haploid assemblies of *S. japonicus* (SJF-hap1), *S. australasicus* (SAM-hap1), and *S. scombrus* (SSM-hap1) with those of *T. thynnus* (TT) and *T. maccoyii* (TM). Syntenic, inverted, translocated, and duplicated regions are shown in grey, yellow, green, and sky blue, respectively. The position of the centromere is shown as a black rectangle for each chromosome, but those on Chr4, Chr11 and Chr24 of the SAM were based on the secondary assembly (SAM-hap2), as the centromere was not detected in the primary assembly.

### Species divergence tree

We constructed the species phylogeny using gene sequences by extracting 6,400 single-copy orthologues from four *Scomber* mackerels (*S. japonicus*, *S. australasicus*, *S. scombrus*, and *S. colias*) and two outgroup species (*Thunnus maccoyii* and *T. thynnus*) and inferred the phylogeny using a supermatrix approach with IQ-TREE. Consistent with the mtDNA-based species tree^29,30^, the resulting tree showed that *S. japonicus* is most closely related to *S. colias*, followed by *S. australasicus*, while *S. scombrus* forms a sister clade to this group (Fig. 1A), although the topology of these phylogenetic trees differs from that inferred from a maximum-likelihood (ML) tree based on whole mitogenome sequences^31^, where *S. japonicus* and *S. australasicus* form a monophyletic clade, with *S. colias* placed outside this group.

### Massive chromosomal rearrangements through pericentric inversion

Chromosomal inversions are known to contribute to reproductive incompatibility between incipient or closely related species^32,33^, with pericentric inversions generally having a more substantial effect on gametogenesis than paracentric inversions when recombination is not suppressed^34^. We identified numerous large-scale chromosomal rearrangements, including megabase-scale inversions and translocations among the three *Scomber* species (Fig. 1B). The number of large inversions ranged from 34 to 41 across each pairwise comparison: 16 pericentric and 19 paracentric inversions between *S. japonicus* and *S. australasicus*, 13 pericentric and 28 paracentric inversions between *S. japonicus* and *S. scombrus*, and 2 pericentric and 32 paracentric inversions between *S. australasicus* and *S. scombrus* (Table 2). Taking divergence times into account, the frequency of interspecific inversions was approximately seven times higher between *S. japonicus* and *S. australasicus* than between either of these Pacific species and *S. scombrus*. This result suggests that more extensive chromosomal rearrangements have occurred between the parapatric species pair than between allopatric pairs. Notably, pericentric inversions appear to have evolved rapidly in the lineage leading to *S. japonicus*, as this lineage exhibited a marked increase in their frequency.

**Table 2.**
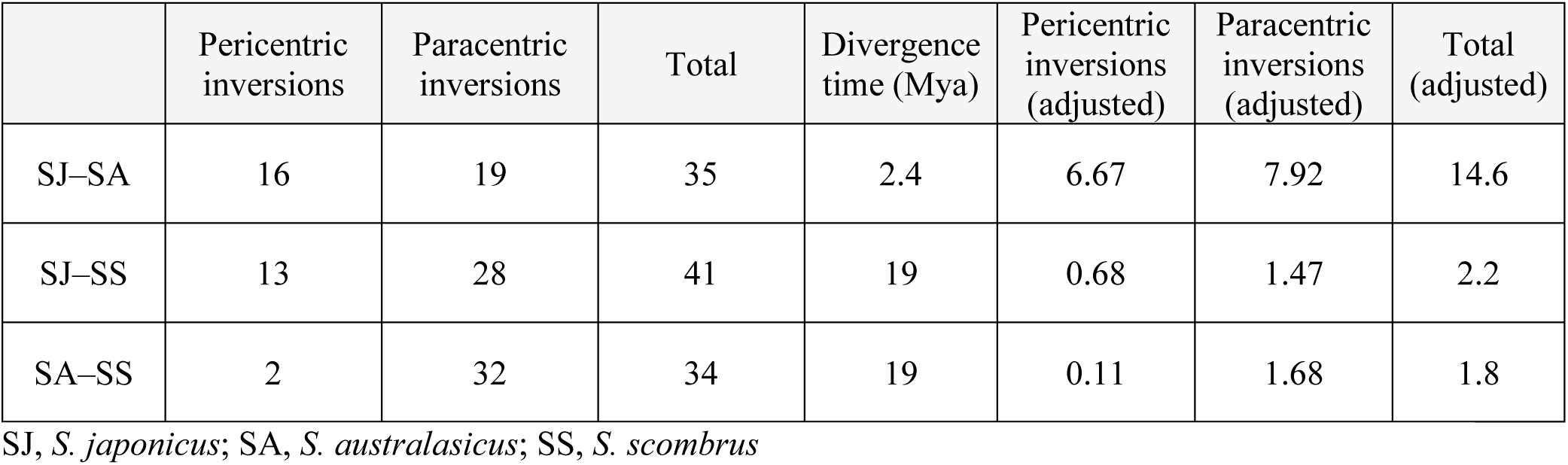
Number of inversions in three *Scomber* species.

|  | Pericentric inversions | Paracentric inversions | Total | Divergence time (Mya) | Pericentric inversions (adjusted) | Paracentric inversions (adjusted) | Total (adjusted) |
| --- | --- | --- | --- | --- | --- | --- | --- |
| SJ-SA | 16 | 19 | 35 | 2.4 | 6.67 | 7.92 | 14.6 |
| SJ-SS | 13 | 28 | 41 | 19 | 0.68 | 1.47 | 2.2 |
| SA-SS | 2 | 32 | 34 | 19 | 0.11 | 1.68 | 1.8 |

SJ, *S. japonicus*; SA, *S. australasicus*; SS, *S. scombrus*

We also observed considerable variation in the frequency of inversions across chromosomes between the two Pacific mackerel species (Supplementary Table S4). In particular, four chromosomes (Chr6, Chr18, Chr20, and Chr24) harboured more than two large paracentric inversions. These extensive interspecies paracentric inversions, along with pericentric inversions, may contribute to hybrid incompatibility and the maintenance of species boundaries in the two parapatric species. In contrast, the absence of large-scale inversions on Chr17 across species pairs suggests that this chromosome may be subject to stronger evolutionary constraints. No apparent association was detected between the number of large inversions per chromosome and the abundance of transposable elements or other repeat sequences (Supplementary Fig. 5).

### Intraspecies inversion

We further identified intraspecific inversions in *S. japonicus* and *S. australasicus* by employing local PCA with nonoverlapping 10-kb windows based on the low-coverage whole-genome resequencing data obtained from wild samples commercially collected from multiple locations (Supplementary Table S5). In total, we identified megabase-scale inversions on Chr24 for *S. japonicus* and on Chr5, Chr6, Chr20, and Chr24 for *S. australasicus* (Supplementary Table S6). As an example, two intraspecific inversions were detected on Chr5 of *S. australasicus*, where a marked increase was observed in the multidimensional scaling (MDS) plot (Fig. 2A). Individuals were clearly grouped into three clusters based on the sequence/genotype in each outlier region (Fig. 2B), with one cluster having higher heterozygosity (Fig. 2C), and the region showed a clear pattern of high LD (Fig. 2D). This estimate is conservative because our approach was limited to identifying inversions >500 kb in length with a minimum allele frequency of 5%. With respect to *S. scombrus*, the density of RAD-seq markers was not sufficient to perform this analysis; thus, intraspecies inversion was not investigated.

**Fig. 2:**
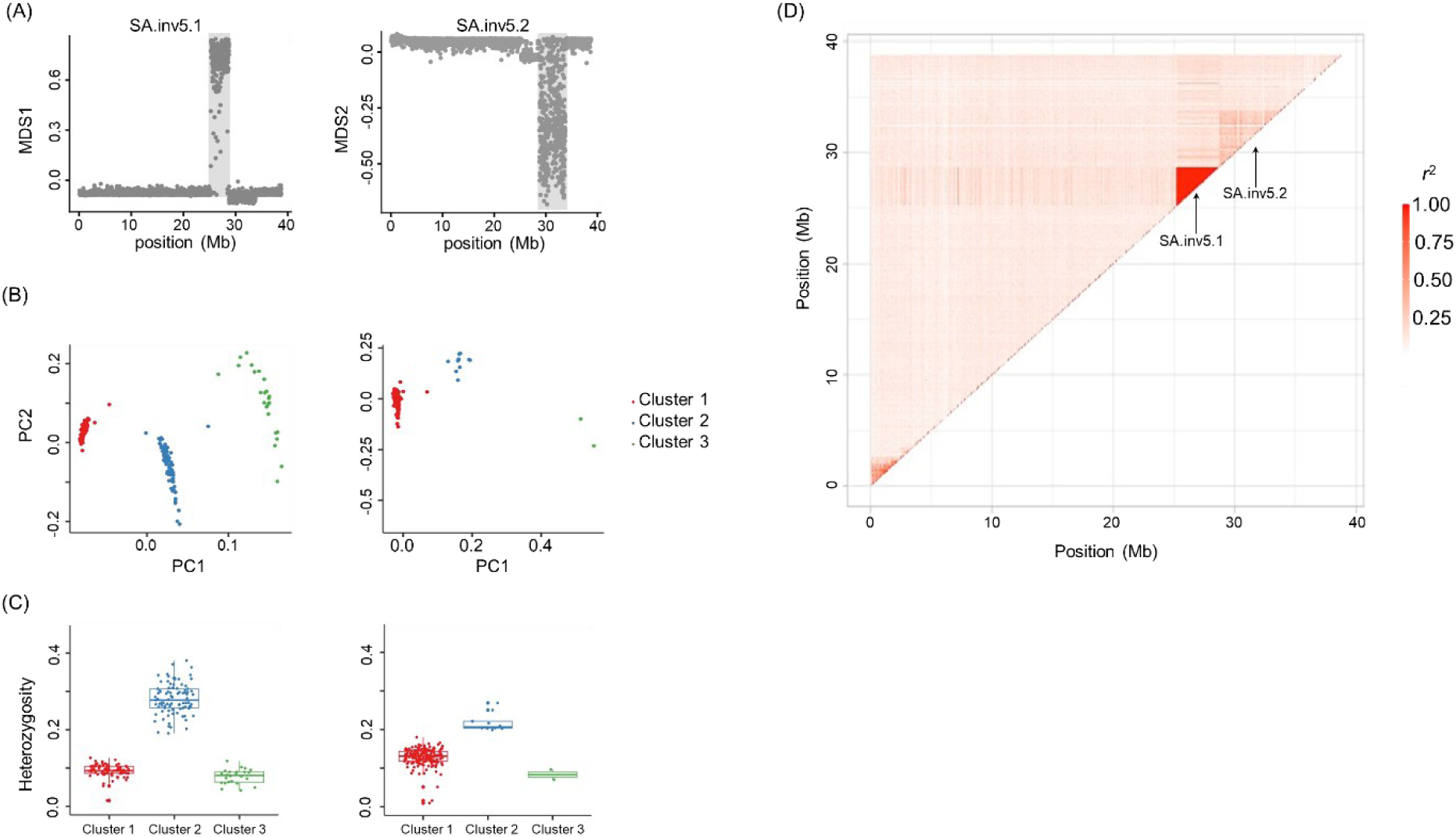
Two intraspecific inversions on Chr5 of *Scomber australasicus* detected by a local principal component analysis (PCA; 10-kb window). **(A)** Multidimensional scaling (MDS) values plotted along genomic positions inferred from the local PCA. **(B)** Sample clustering identified by *k*-means clustering based on the PCA results for each region. **(C)** Variation in heterozygosity within the inverted region among clusters. Individual heterozygosity values represent the mean within the region; cluster names correspond to those in panel (B). In panels (A)–(C), the left panel represents SA.inv5.1 and the right panel represents SA.inv5.2, respectively. **(D)** Linkage disequilibrium (LD) heatmap for Chr5 of *S. australasicus*. The colour scale indicates the degree of LD (*r*²).

### Characterization of inversion breakpoints

Inversion breakpoints are frequently located within highly repetitive sequences^35^. Consistent with this result, we observed an accumulation of tandem repeat arrays around chromosomal inversion breakpoints in the three *Scomber* mackerels. These enrichment patterns could be categorized into three types (Fig. 3A). First, both breakpoints were near duplicated repeat arrays, a pattern predominantly associated with paracentric inversions (39 of 41 cases) (Fig. 3A, top left panel, and Fig. 3B; Supplementary Table S7). Second, one breakpoint was near centromeric repeats, whereas the other was flanked by a tandem repeat array either near the telomere or at a paracentromeric position (Fig. 3A, top right panel). This pattern was more common in pericentric inversions (18 of 22 cases). Third, only one of the two breakpoints was flanked by a tandem repeat array (Fig. 3A, bottom left panel), which was observed in both pericentric and paracentric inversions (11 and 20 cases, respectively). Notably, the breakpoints of paracentric inversions observed on two distinct chromosomes (Chr8: 20,635,651–25,113,923 bp and Chr13: 12,017,864–17,234,664 bp) shared a very similar duplicated array composed of a 354-bp satellite repeat. Duplicated satellite DNA arrays were also found at both breakpoints of intraspecies inversions in *S. australasicus* (SA.inv5.2, SA.inv6, SA.inv20, and SA.inv24.1), although the repeat units varied among the loci. Recurrent use of rearrangement breakpoints was observed, with the same genomic regions repeatedly involved in independent rearrangements, for example, on Chr13 and Chr24 (Supplementary Fig. 8). Taken together, these findings suggest the presence of genomic loci that promote chromosomal rearrangements - so-called rearrangement hotspots - that are characterized primarily by dense repeat arrays.

**Fig. 3:**
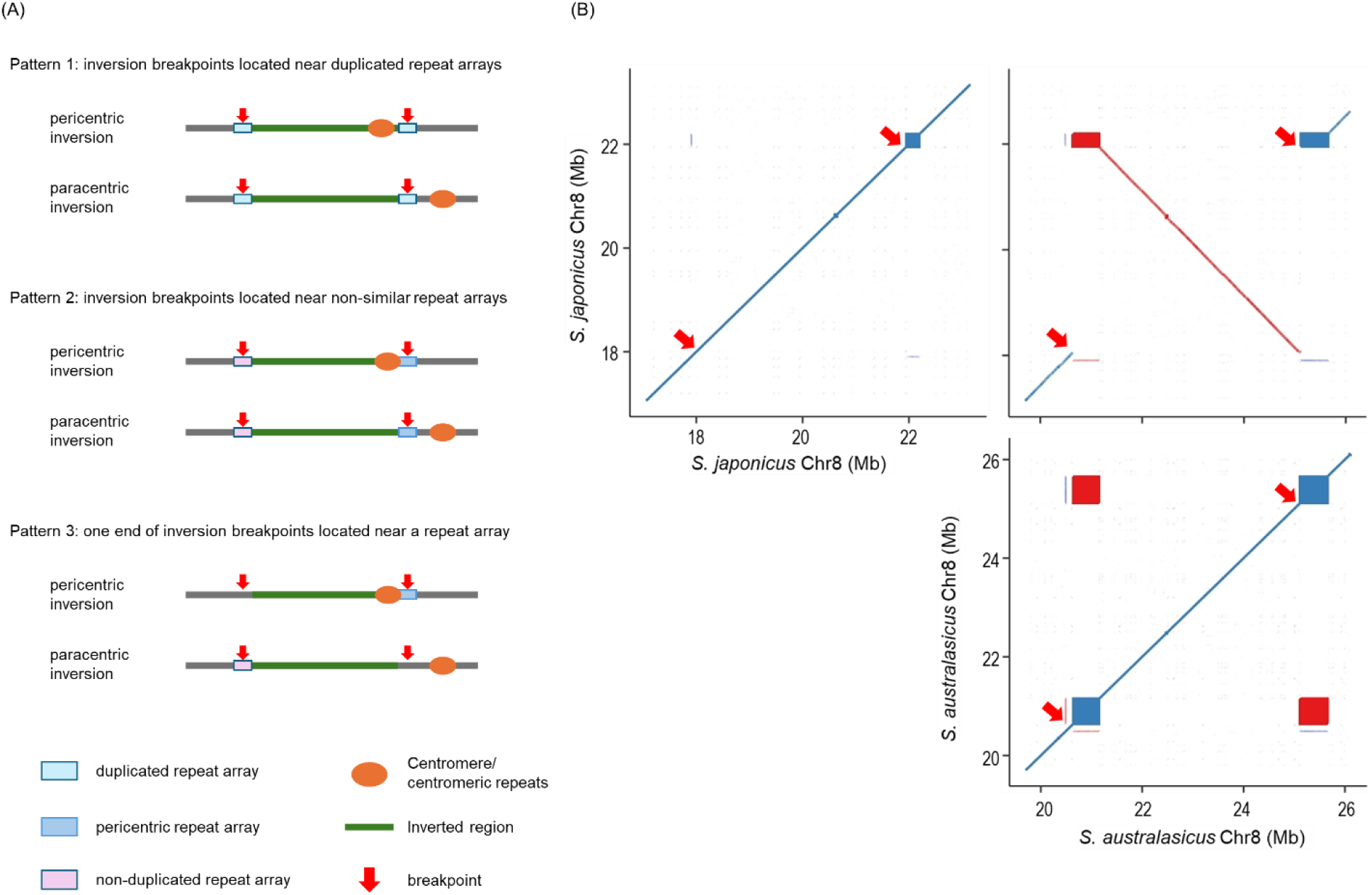
Enrichments of repetitive sequences at the inversion breakpoints. Illustration of three patterns of repeat arrays **(A)** and an example of Pattern 1 observed in the inversion on Chr8 between *S. japonicus* and *S. australasicus* **(B)**. Dots represent alignments longer than 10 kb within *S. japonicus* (top left panel) and *S. australasicus* (bottom panel) and between these two species (top right panel). Reverse orientations are coloured red and red arrows indicate breakpoints.

### Sex chromosome identification

Given that sex determining regions (SDRs) are known to exhibit a greater propensity for the accumulation of large SVs than autosomes and tend to diverge rapidly among closely related species, we conducted genome-wide association studies (GWASs) based on the same resequencing data used for the local PCA (136 males and 145 females of *S. japonicus* and 96 males and 92 females of *S. australasicus*) to characterize their previously unidentified sex chromosomes and SDRs. We detected strong signals associated with phenotypic sex in a region of Chr24 spanning 11.5 Mb (3.0–14.5 Mb) in *S. japonicus* (Fig. 4A) and in a region of Chr16 spanning approximately 12.5 Mb (10.0–22.5 Mb) in *S. australasicus* (Fig. 4B). A comparison of female- and male-specific SNPs in the SDR revealed female heterogamety (ZZ/ZW) in *S. japonicus* and male heterogamety (XX/XY) in *S. australasicus*, consistent with a previous report^20^. While the distribution of sex-biased SNPs overlapped with the strongly associated region in *S. japonicus*, it was concentrated within a limited 4.8 Mb region (14.7–19.5 Mb) in *S. australasicus* (Supplementary Fig. 9). Thus, we determined the SDR for *S. japonicus* as the 11.5 Mb region on Chr24 and that for *S. australasicus* as the 4.8 Mb region on Chr16. We also confirmed that the previously designed sex diagnostic markers reside in the SDR for each species (Supplementary Table S8).

**Fig. 4:**
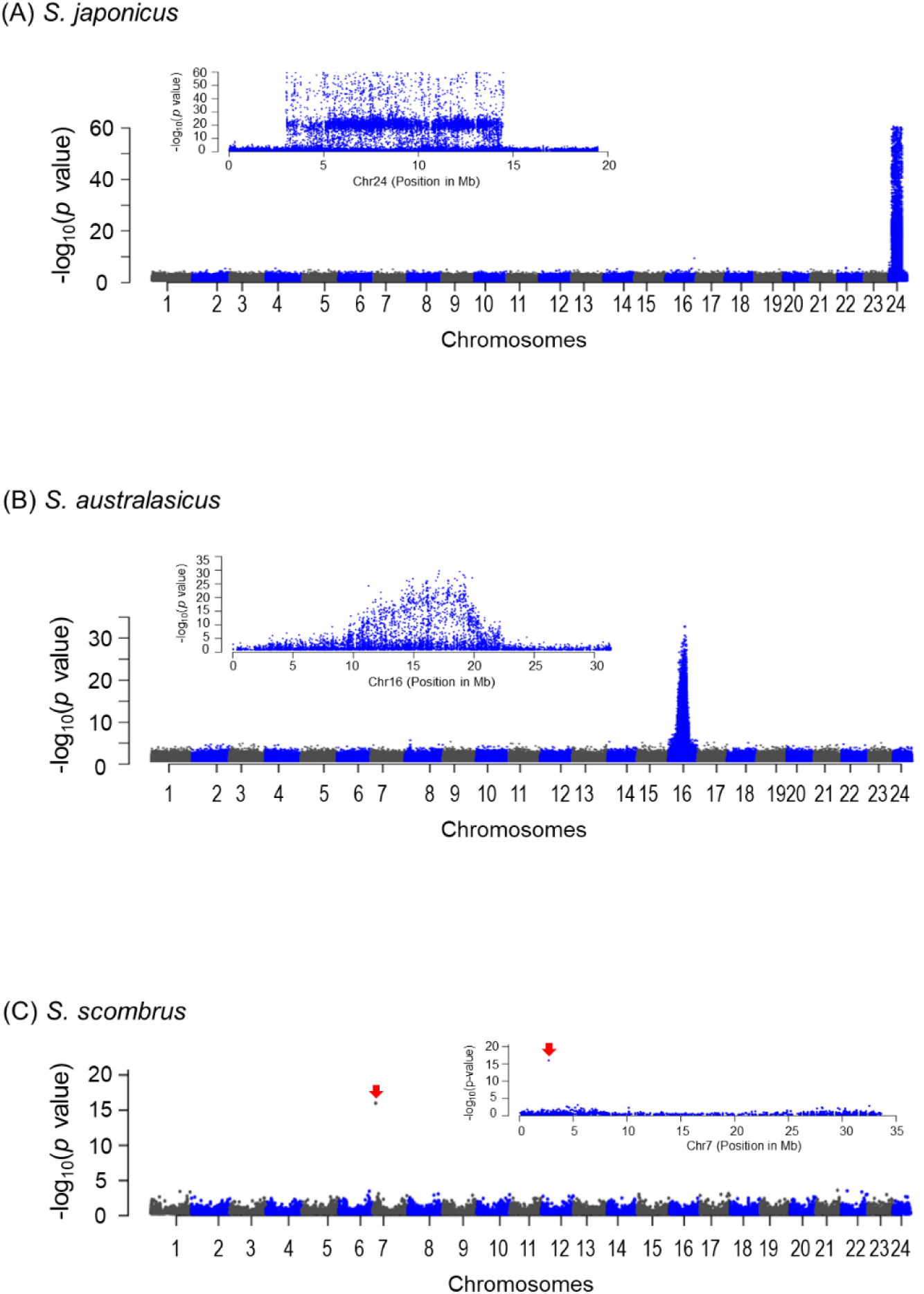
Manhattan plots of genome-wide associations for phenotypic sex. Associations between genome-wide SNPs and phenotypic sex were analysed by generating Manhattan plots for *S. japonicus* **(A)**, *S. australasicus* **(B)**, and *S. scombrus* **(C)**. In the main panel, the X-axis indicates chromosomal positions, and the Y-axis shows the -log_10_ *p* values of individual SNPs, each represented by a dot. Insets present enlarged views of the chromosomes in which significant SNPs were detected. Red arrows in panel (C) indicate significant SNPs.

We also investigated the sex chromosomes in *S. scombrus* using RAD-Seq data (158 males and 143 females) to gain insights into the evolutionary history of the two different heterogametic sex determination systems in the two species and identified a single male-biased RAD variant that was detected in 88 males but only 4 in females (Supplementary Fig. S10A). Utilizing this RAD variant, we identified a male-specific locus on Chr7 of SSM-hap1 (Fig. 4C) but not in SSM-hap2. These results indicate that this species has a male heterogametic sex-determination system, with the SDR on Chr7.

Taken together, these results suggest that the three genetic sex determination systems evolved independently in the three species, highlighting the evolutionary lability of sex chromosomes.

### Structural variations in sex chromosomes of *S. japonicus*

After the SDR in each species was detected, we conducted an in-depth analysis of the structural characteristics of the sex chromosomes using the results of the SV analysis described above. In the case of *S. japonicus*, we detected three large inversion polymorphisms on Chr24: INV1 (3.02–4.93 Mb), INV2 (4.95–14.50 Mb), and INV3 (14.50–15.68 Mb) (Supplementary Fig. 11A). However, the results of the GWAS indicated that INV3 is not associated with sex, but INV1 and INV2 are. This result was confirmed with three additional haplotype-resolved assemblies; the inverted configuration within INV3 regions was not restricted to females (Supplementary Fig. 11B). We consistently observed elevated *F*_ST_ between the sexes at INV1 and INV2 but not at INV3 (Supplementary Fig. 11C). These findings indicate sex-specific or sex-biased inversion karyotypes for INV1 and INV2, whereas INV3 remains polymorphic across the population. These interpretations are consistent with the reanalysed local PCA, which revealed sex-specific clustering for INV1 and INV2 but not for INV3 (Supplementary Fig. 12A, B). We also observed a female-specific increase in LD across INV1 and INV2, indicating a strong suppression of recombination in this sex-associated region (Supplementary Fig. 12C), which explains the enrichment of large insertion/deletion variations, specifically TEs such as LINEs/LTRs (∼4–7 kb) and helitrons (∼7 kb), specific to INV1 and INV2 (Supplementary Fig. 13).

Polymorphic inversions on both the Z and W chromosomes are particularly noteworthy as potential drivers of chromosomal restructuring, given that they may promote the progressive loss of recombination in sex chromosomes, a recurrent feature of their evolutionary history^36,37^. We tested this possibility by analysing the genotype frequencies of INV3 orientations (Supplementary Table S9). The occurrence of homozygotes for the inverted orientation in both sexes indicates the presence of both Z-linked and W-linked inversions. Nonetheless, the significantly lower frequency in females (*P* = 0.00789, chi-square test) implies a sex-specific fitness effect of the inverted region^38,39^, which may in turn facilitate the expansion of the nonrecombining region through the fixation of the Z-linked inversion. Alternatively, the Z-linked inversion may be neutral but has arisen first, with the W-linked inversion emerging some time later through recombination between Z and W.

We further investigated the effect of the polymorphic inversion on the evolution of sex chromosomes by examining genetic differentiation between the standard and inverted arrangements at the INV3 locus. We performed an *F*_ST_ analysis between individuals who were homozygous for the standard genotype and those who were heterozygous; individuals who were homozygous for the inverted arrangements were excluded because of their limited sample size. As expected, *F*_ST_ was increased across the INV3 locus, spanning approximately 1.2 Mb (Fig. 5A, B). Notably, an additional *F*_ST_ peak was observed in a 0.8 Mb region at INV2, located on 13.7–14.5 Mb on ChrZ (Fig. 5A) and orthologous to the 5.1–5.9 Mb region on ChrW (Fig. 5B). We conducted a local PCA using only ZZ males to test whether this divergence in the 0.8 Mb region had arisen among the ChrZ sequences. This analysis revealed a sharp peak in the first coordinate (MDS1) within this region, together with a secondary peak at INV3 (Fig. 5C). PCA of the 0.8 Mb region revealed that ZZ individuals were grouped into two clusters, whereas the results of the *F*_ST_ analysis showed clear genetic differentiation between the two clusters (Fig. 5D). These results indicate the presence of at least two divergent haplotype groups within the SDR of chromosome Z. This divergence may be associated with the polymorphic inversion INV3 located adjacent to the SDR, as the LD heatmap and LD decay analyses revealed genotype associations between this region and INV3 (Supplementary Fig. 14). We cannot exclude the possibility that additional haplotype diversification has also occurred within the 0.8 Mb region on ChrW.

**Fig. 5:**
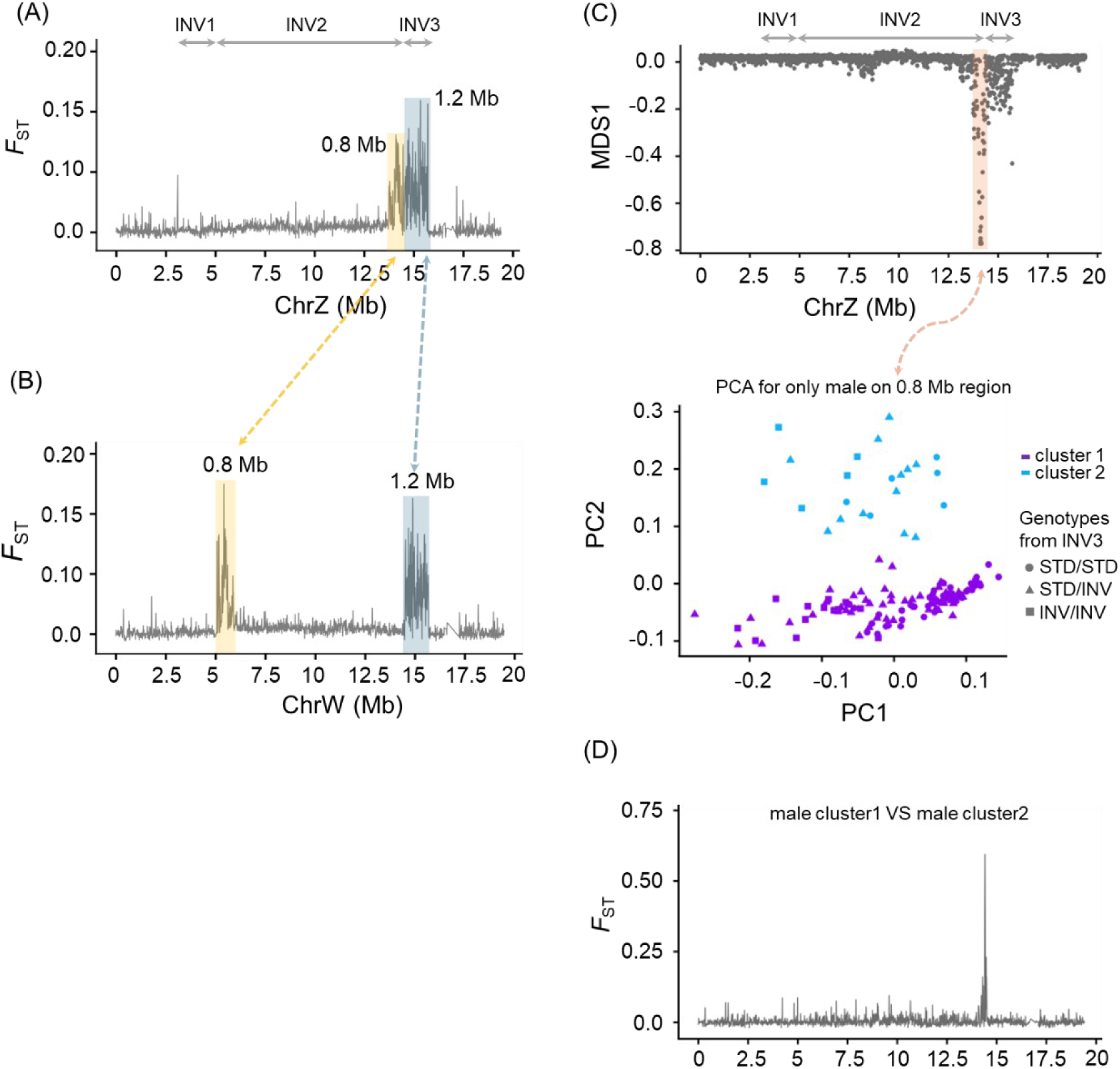
Multiple Z chromosome lineages revealed by an additional recombination suppression region near the polymorphic inversion (INV3) on the sex chromosome of *S. japonicus*. **(A)** *F*_ST_ between STD/STD vs. STD/INV individuals along with the ChrZ reference sequence and **(B)** along with the ChrW reference sequence. **(C)** Multidimensional scaling (MDS) values plotted along genomic positions inferred from the local PCA (top panel) and sample clusters identified by *k*-means clustering based on the PCA results for the differentiated region (bottom panel). Only males were included in this analysis. In the bottom panel, each individual was plotted with different colours based on the clustering result in the 0.8 Mb region and shapes based on the genotype in the INV3 region (circle, STD/STD; triangle, STD/INV; and square, INV/INV). (D) *F*_ST_ between the clusters in males based on the 0.8 Mb region on ChrZ.

We first searched for genes specific to either the Z or W chromosome to identify potential candidates for primary sex-determining genes. However, all the annotated genes (306 protein-coding genes) in the SDR are shared between these chromosomes. Among these shared genes, no homologues of known sex-determining genes from other teleost species were identified. Nevertheless, several genes, such as *sox1b* in INV2, exhibit a degree of similarity to genes that are potentially involved in gonadal differentiation in other fish species (Supplementary Table S10)^40,41^.

### Structural variations in the sex chromosome of *S. australasicus*

In contrast to *S. japonicus* described above, our SNV analysis, in particular the comparison of haplotype-resolved genome assemblies (Fig. 1), did not identify any obvious large inversions on the sex chromosome (Chr16), even though the GWAS and heterozygosity levels of sex-biased SNPs indicated a substantial reduction in recombination between the sex chromosomes (10–23 Mb) (Fig. 4B and Supplementary Fig. 9). Additional confirmation from long-read alignments further supported the absence of large inversions in a total of three XY males and two XX females (Supplementary Fig. 15). These results suggest the presence of poorly understood mechanisms that suppress recombination in a large collinear region^42–44^.

We investigated the suppression of sex chromosome recombination in more detail by analysing LD and unexpectedly identified three distinct LD blocks in male genomes. One of these blocks (LD1: 9.6–19.35 Mb) encompassed the SDR (14.7–19 Mb) and was flanked by the other two blocks (LD2: 1.75–9.6 Mb; LD3: 19.35–23.0 Mb), with further LD observed between these flanking LDs (Fig. 6A), suggesting the presence of multiple Y haplotypes. Consistent with this result, a reanalysis of the local PCA clearly separated males into multiple clusters, three in LD2 and four in LD3 (Fig. 6B, C), while females formed a single cluster together with a subset of males, Cluster 1 in both LD2 and LD3. *F*_ST_ analysis between females and each of the male clusters revealed marked heterogeneity in the size of the recombination suppression region along the sex chromosome (Fig. 6D); the size of the suppressed region varied from approximately 10 Mb, encompassing LD1 (orange in the left panel, purple in the right panel), to approximately 20 Mb, spanning all three LD blocks (LD2, green and purple; LD3, magenta, light blue, and gold). *F_ST_* between males and females was consistently high in LD1 across all pairwise comparisons, whereas the elevated *F_ST_* in LD2 and LD3 was observed in a subset of comparisons (4 of 7 in LD2 and 5 of 7 in LD3). This pattern suggests that these evolutionary strata may be of similar age or that incomplete recombination suppression hindered divergence, particularly within LD1.

**Fig. 6.**
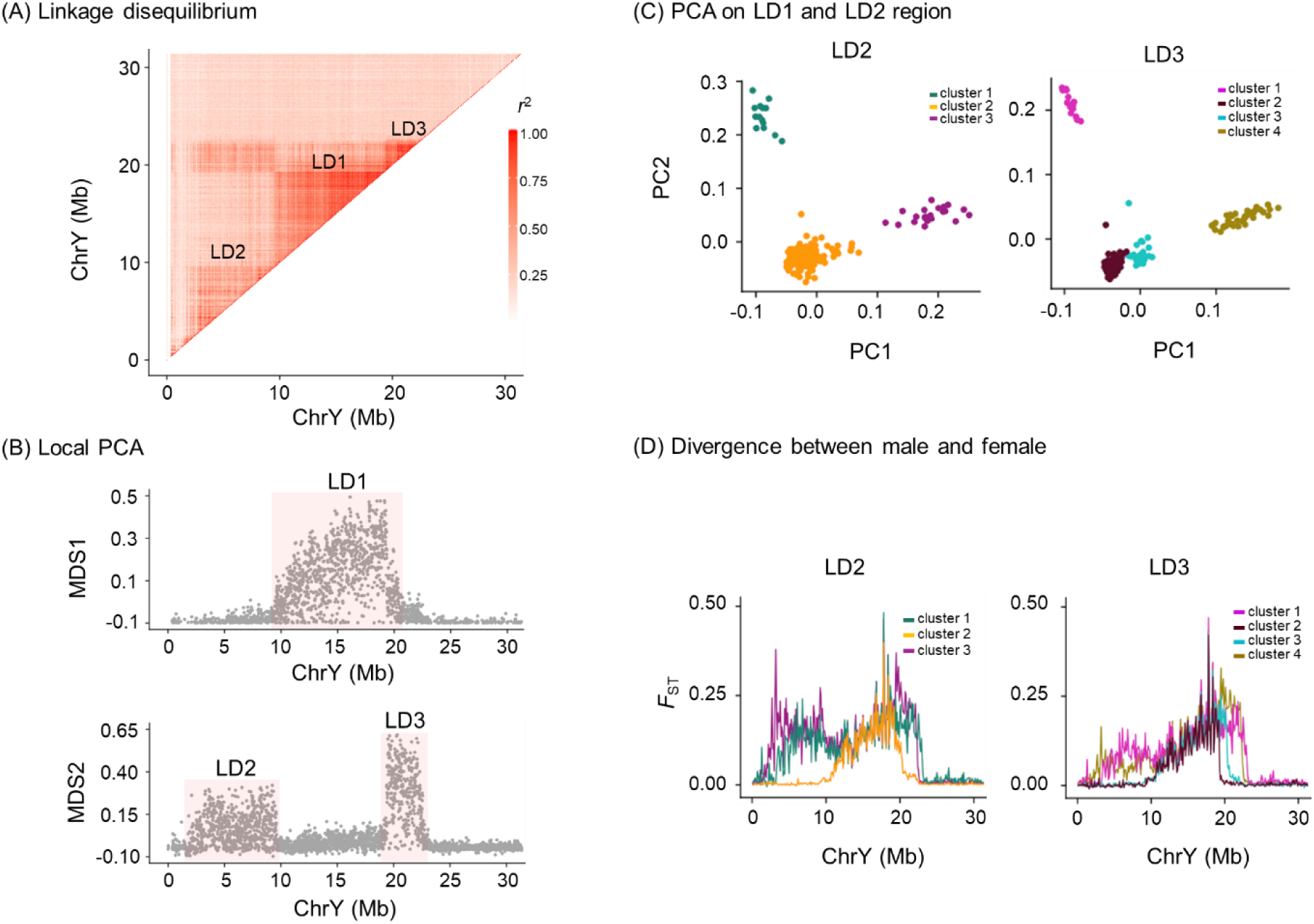
Multiple Y chromosome lineages associated with variable divergent regions in *S. australasicus.* **(A)** LD for the sex chromosome (Chr16) of *S. australasicus*, shown as maximum *r*^2^ values for paired windows across the chromosome. Maximum *r²* values were calculated using samples from both males and females. Scales for *r*^2^ values are provided. **(B)** Local PCA to detect the outlier region with a 10-kb window. Distances between local PCA maps are shown on the MDS1 (top panel) and MDS2 axes (bottom panel), with outlier windows shaded in red. **(C)** Clustering of samples by PCA for the outlier region on LD2 (left panel) and LD3 (right panel) on MDS2, assigned using *k*-means clustering. **(D)** *F*_ST_ across the length of ChrY between males of each cluster and females based on PCA on LD2 (left panel) and LD3 (right panel).

Although our search did not identify protein-coding genes specific to either the X or Y chromosome, the SDR contains two genes homologous to known teleost sex-determining genes, namely, *gsdf* and *bmpr1bb*^45–47^, among the 164 genes shared by both chromosomes (Supplementary Table S11).

### Differences in the sex-determining regions in the two species

Recent studies have suggested that recombination suppression on sex chromosomes can occur through mechanisms other than inversions. For example, Jeffries *et al*. (2021)^48^ proposed an alternative to the inversion model: sequence divergence accumulating near the SDR boundaries can progressively expand the suppressed region (a “divergence-driven” model), generating diversity patterns distinct from inversion-driven suppression. The two *Scomber* fishes provide a rare opportunity to analyse these contrasting patterns. The *F*_ST_ pattern in LD1 for a subset of *S. australasicus* males (orange in the left panel and purple in the right panel of Fig. 6D) fits the divergence-driven model, forming a localized peak, rather than the broad, plateau-like elevation expected under the inversion-driven model observed in *S. japonicus* (Supplementary Fig. 11C). Indeed, large TEs are enriched within the SDR in *S. australasicus* compared to that in *S. japonicus* (Supplementary Fig. 13, 16 and 17), a pattern that may reduce recombination by inhibiting the homology-searching process when TEs are inserted near the SDR boundaries. These observations are consistent with the hypothesis that TEs have contributed to the progressive loss of recombination at the SDR in *S. australasicus*. Notably, the suppressed recombination region in *S. australasicus* appears to have expanded or contracted as a whole in units of the three LDs. The underlying cause was likely linked selection acting on the sexually antagonistic loci and/or on deleterious mutations located on LD2 and LD3^49^.

Neutrality tests further highlighted a distinct evolutionary process that shaped the suppression of recombination in the SDRs of the two species (Fig. 7). Tajima’s D analysis of heterogametic individuals (ZW) of *S. japonicus* showed the positive values expected under balancing selection in the nonrecombining SDR, whereas in *S. australasicus*, the heterogametic individuals (XY) had negative values. The pattern observed in *S. australasicus* may reflect the recent origin of the Y haplotypes under positive selection, which has not allowed sufficient time for the accumulation of mutations detectable as balancing selection. In addition, incomplete recombination suppression in the collinear SDR may also contribute by permitting occasional low-frequency gene exchange between the X and Y chromosomes.

**Fig. 7.**
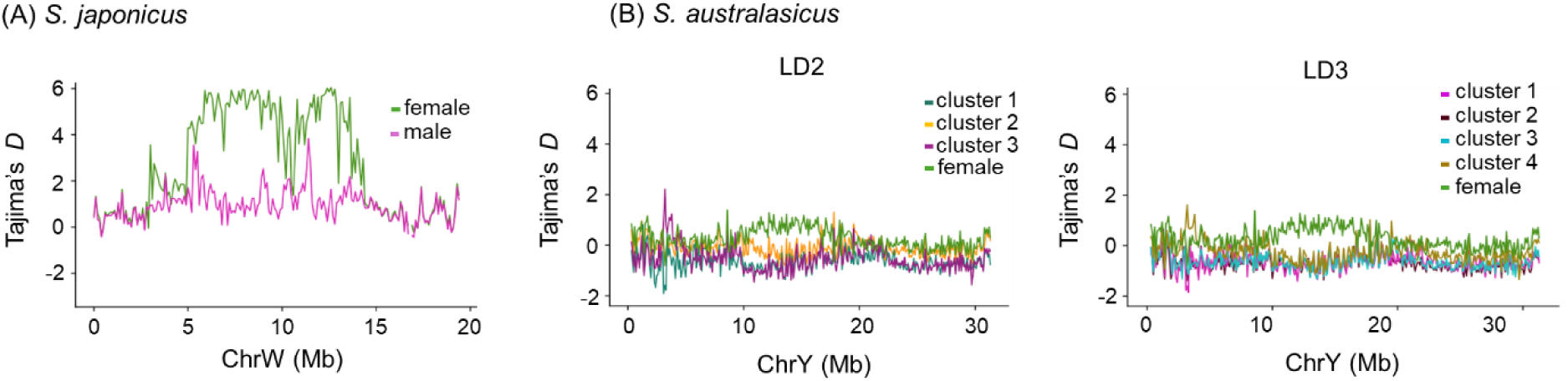
Neutrality test on the sex chromosomes. **(A)** Tajima’s *D* values for males and females of *S. japonicus* are plotted on the y-axis, with chromosomal positions along ChrW shown on the x-axis. **(B)** Tajima’s *D* values for females and for each of the male clusters based on the LD2 (left panel) and LD3 (right panel) regions of *S. australasicus* are plotted on the y-axis, with chromosomal positions along ChrY shown on the x-axis. Tajima’s *D* was calculated as a neutrality test in nonoverlapping 100-kb windows.

### Structural variations in the sex chromosomes of *S. scombrus*

In contrast to the cases of the two Pacific *Scomber* fishes, the differentiated region between the two haploid assemblies of the sex chromosomes of *S. scombrus* (Chr7X and Chr7Y) was only limited to an ∼14 kb insertion variant on the Y chromosome (Chr7Y: 2,918–2,932 kb) in the region where the male-biased RAD sequence is mapped (Fig. 4C and 8A). A BLAST search showed high similarity between an 8 kb segment in the 14 kb locus and an autosomal region on Chr5 (22,206,152–22,214,080 bp), suggesting that the duplicated and transposed sequences originating from this autosomal region gave rise to the male-specific region. We then designed a genetic marker that amplifies the Y-specific region on Chr7 and its autosomal paralogue of different sizes (Supplementary Table S12) and confirmed its male-specific presence in newly obtained wild-caught individuals (14 males and 16 females) (*P* = 6.877×10^-9^, Fisher’s exact test) (Supplementary Fig. 10B). By leveraging the genetic marker, we identified four genotypic males and six genotypic females in a set of publicly available whole-genome resequencing data (NCBI BioProject Accession: PRJNA253681) (see Supplementary Table S13) and used these individuals to specify the SDR with finer resolution. The *F*_ST_ analysis indicated that genetic differentiation between the X and Y chromosomes does not extend to the boundaries of the male-specific inserted region (Fig. 8B). Moreover, a male-to-female coverage ratio analysis along Chr7 revealed an approximately 14 kb region (Chr7Y: 2,918,935–2,932,707 bp) with marked differences in the coverage ratio (Fig. 8C). Notably, a segment of ∼377-bp inverted-duplicate repeats was identified at the borders of the duplicated segment, suggesting that nonallelic homologous recombination (NAHR) facilitated the duplication and translocation of this region from the autosome (Chr5) to the Y chromosome (Chr7Y), functioning as a sex-determining locus (Fig. 8D).

**Fig. 8.**
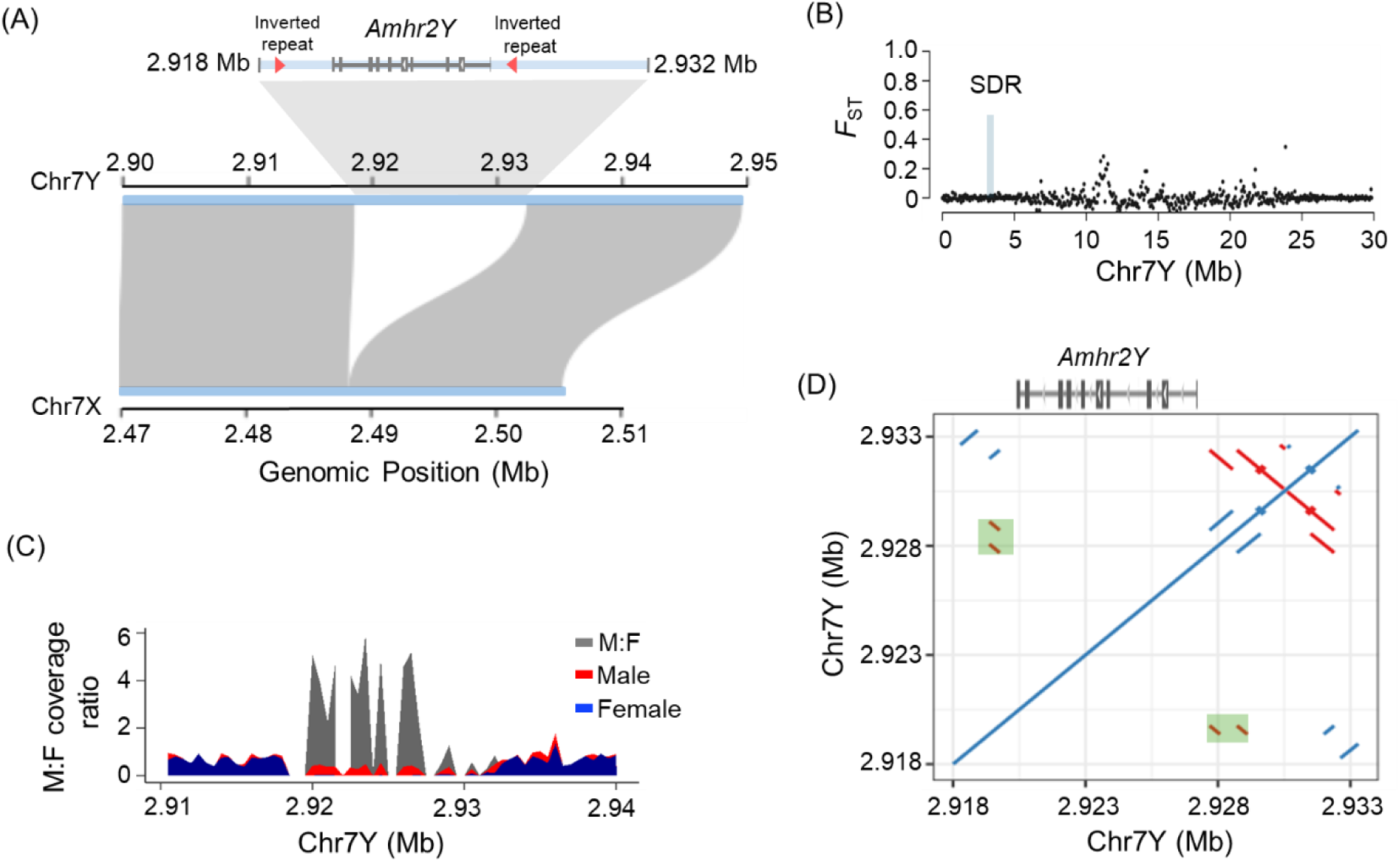
Y-specific region on the sex chromosome (Chr7) of *S. scombrus*. **(A)** A synteny plot between Chr7Y and Chr7X around the sex-determining region represents the 13.77-kb Y-specific insertion, which harbours the single gene *amhr2* (*amhr2Y*). The red triangles show 377-bp inverted repeats at the proximal and distal borders of *amhr2Y.* These inverted repeats are assumed to drive the duplication and transposition of the *amhr2* gene on the Y chromosome (Chr7) from the autosome (Chr5) via nonallelic homologous recombination (NAHR). **(B)** *F*_ST_ in nonoverlapping 100-kb windows between males and females across the length of Chr7. **(C)** Male-to-female coverage analyses in nonoverlapping 500-bp windows around the sex-determining region. **(D)** Self-aligned dot plot of the Y-specific region. The green shaded areas are the 377-bp inverted repeats at the proximal and distal borders of *amhr2Y*.

The duplicated segment included a single protein-coding gene: *amhr2*. This gene, or its paralogues, functions as the master sex-determining gene in at least nine genera of fish, including *Takifugu* pufferfish and *Pangasius* catfish^47,50–52^. Therefore, *amhr2* is a strong candidate for the primary sex-determining gene in *S. scombrus*.

These results indicate that the sex-determining locus evolved through a third, distinct mechanism of recombination suppression—namely, the duplication and translocation of a coding gene—compared with those observed in the two Pacific species mentioned above, highlighting this genus as a model for the evolution of sex chromosomes.

## Discussion

In this study, we investigated the evolutionary mechanisms underlying hybrid incompatibility between two parapatric Pacific mackerels, *S. japonicus* and *S. australasicus*, hypothesizing that chromosomal rearrangements are the major driving forces. We examined this hypothesis by generating high-quality, haplotype-resolved *de novo* genome assemblies for each species. Together with a reassembled haplotype-resolved genome of *S. scombrus* reconstructed from public data, our comparative genomic analyses enabled a detailed karyotype comparison, revealing extensive chromosomal rearrangements. Notably, the rate of chromosomal inversions was approximately sevenfold higher between the parapatric Pacific pair than between allopatric species pairs. Complex rearrangements involving megabase-scale inversions and confounded translocations were also observed in the two Pacific species. We further conducted large-scale structural variation analyses using local PCA based on an extensive dataset comprising more than one hundred individuals from multiple regions. These analyses enabled a complementary investigation of intraspecies inversions. The analyses also revealed that the sex chromosomes of the three species underwent independent evolution. Convergent evolution of large-scale recombination suppression (>10 Mb) was identified on the sex chromosomes of the two Pacific species through chromosomal inversion in *S. japonicus* and, most likely, TE-mediated sequence differentiation in *S. australasicus*, whereas in *S. scombrus*, recombination suppression was confined to an ∼14 kb hemizygous region formed by duplication and translocation. The population-level analysis also provided a snapshot of the ongoing evolution of the recombination suppression region in *S. japonicus* and *S. australasicus* and, in particular, revealed diverse Y chromosome lineages with markedly different lengths of the suppressed region in *S. australasicus*. Collectively, these findings highlight the dynamic nature of genome evolution in *Scomber* and provide a genomic framework for understanding how chromosomal rearrangements shape species boundaries, reproductive incompatibility, and sex chromosome turnover.

Megabase-scale chromosomal rearrangements can occur with and without centromere involvement. Pericentric inversions—inversions that involve the centromere—are a major mechanism reshaping chromosome structure and have been documented across diverse taxa, including vertebrates^53,54^, such as fish^55^. A large-scale survey of passerine birds (411 species) revealed that pericentric inversions were common among parapatric species pairs and that their fixation rate of inversions correlated positively with range overlap, suggesting that pericentric inversions play a key role in restricted gene flow^53^. Similarly, we detected pericentric inversions among *Scomber* genomes. Compared with *S. australasicus*, the lineage leading to *S. japonicus* exhibited extensive chromosomal restructuring, with pericentric inversions affecting two-thirds of the chromosomes. The rate of pericentric inversion evolution—measured as the number of events per million years—exceeded that of passerine birds (maximum 5.1 per Mya), underscoring an exceptionally rapid pace of karyotypic evolution in this lineage.

Although the mechanisms driving megabase-scale chromosomal inversions remain largely unknown, the recurrent use of inversion breakpoints has been observed in several taxa^56^, suggesting that certain genomic regions are repeatedly involved in structural rearrangements, possibly reflecting inherent fragility or adaptive reuse of specific chromosomal segments. Consistent with these observations, our data also revealed the recurrent use of inversion breakpoints across multiple chromosomes (e.g., Chr13 and Chr24) in *Scomber* genomes. Inversion breakpoints are sometimes associated with segmental duplications, TEs and centromeric repeats^56^. Our breakpoint analysis also identified large arrays of repetitive sequences flanking breakpoints of chromosomal inversions in the *Scomber* genomes, although TE enrichment was not observed at breakpoints. Duplicated satellite repeat arrays were located near both ends of the inversion breakpoints (type I in Fig. 3A), suggesting that they arose by nonallelic homologous recombination (NAHR) between these duplicated repeat arrays. A 354-bp satellite array was conserved among the segmental duplications at the breakpoints of the paracentric inversions on Chr8 and Chr13. Similarly, the four intraspecies polymorphic inversions in *S. australasicus* follow the same pattern. In contrast, most pericentric inversions utilized a nonduplicated repeat array: one end resides in pericentromeric repeats adjacent to the centromere, and the other within paracentromeric regions enriched for short tandem repeats (∼50 bp). These features suggest that these pericentric inversions were facilitated by recombination through nonhomologous end joining (NHEJ), a major mechanism of inversion formation in invertebrates that has also been reported in vertebrates such as the Atlantic herring^57^. Collectively, our findings indicate that rapid chromosomal inversions in Pacific *Scomber* species arose through two distinct mechanisms, namely, NAHR and NHEJ, both of which rely on repeat arrays that serve as hotspots for chromosomal breakage and rearrangement. These hotspots may accelerate karyotypic evolution and contribute to hybrid incompatibility between the two parapatric Pacific mackerels.

Intraspecific chromosomal inversions often reorganize multiple genes associated with ecological traits and, by suppressing recombination, preserve coadapted allelic combinations as supergenes (reviewed in^35^). Such inversions underlie adaptive divergence in various fish, including Atlantic cod^58,59^, Atlantic herring^57^, catfish^55^, and Waccamaw darter^60^. For example, recent (< 2 Mya) large inversions in Atlantic cod maintain supergenes linked to migratory versus resident ecotypes. In contrast, no distinct population structure or ecotype has been reported for the two Pacific *Scomber* species around Japan^61^. Nevertheless, we identified one intraspecific megabase-scale inversion in *S. japonicus* and four in *S. australasicus* (allele frequency = 0.04–0.44), with no apparent differences in inversion frequency between geographically distant populations, suggesting the absence of strong ecological selection. However, given its wide geographic distribution, expanding sampling regions, especially towards the southern boundary, is anticipated to yield clearer insights into the ecological roles of intraspecific inversions observed in these species. Specifically, in *S. japonicus*, INV3 on Chr24, the non-sex-associated polymorphic inversion located beside the SD region (Supplementary Table S9), showed significantly lower frequency in females, implying possible ecological or reproductive relevance.

The roles of sex chromosomes and their turnover in speciation have long been debated^62^, with the expansion of recombination-suppression regions regarded as a key process in their evolution^63^. In *S. japonicus* (Chr24), the sex chromosome harbours an extensive LD region formed by two sex-linked large tandem inversions with an additional LD region generated by the adjacent polymorphic inversion (INV3). On the Z chromosome, we detected signatures of expansion (or erosion) of the recombination suppression region from the polymorphic inversion towards the sex-linked region, indicating ongoing sex chromosome differentiation. The expansion or contraction of nonrecombining region around the SDR was even more pronounced in *S. australasicus* (Chr16). In this species, recombination suppression at the SDR likely occurred through the insertion of large TEs rather than through inversions^64^. The population genetic analysis further revealed multiple Y haplotypes differing in the length of recombination suppression regions. While intraspecific variation in the extent of sex chromosome differentiation is expected, particularly in systems with young sex chromosomes, empirical evidence for size variation in recombination-suppression regions within species remains scarce^65^. In the guppy *Poecilia parae*, multiple Y haplotypes with varying sizes of recombination-suppressed regions have evolved across ecotypes, a pattern attributed to negative frequency-dependent selection^66^. In contrast, *S. australasicus* harbours multiple coexisting Y haplotypes in at least two populations. The emergence and maintenance of multiple Y haplotypes are therefore likely driven by evolutionary processes distinct from those inferred for *P. parae*. For example, multiple Y haplotypes of *S. australasicus* may be formed through linked-selection acting on sexually antagonistic loci and/or on deleterious mutations located on LD2 and LD3^49^ or alternative strata formation models^36,67^. Despite these mechanistic differences, the sex chromosomes of the two *Scomber* species showed common evolutionary dynamics, including the coexistence of multiple sex-linked haplotypes and variation in the length of recombination suppression regions. These patterns may reflect underlying chromosomal instability, as supported by the presence of large-scale inversions both within and between species. Alternatively, they may represent a more general process in sex chromosome evolution that remained undetected in studies using limited sample sizes. The large effective population size (*Ne*) of these species may have facilitated the detection of this process before a fixation on specific haplotypes^68^.

In contrast to the two Pacific species, the differentiation and recombination suppression in the sex chromosome of *S. scombrus* (Chr7) was limited to a narrow ∼14 kb region. Sex chromosome turnover in teleosts frequently involves the duplication and translocation of genes in the TGF-β signalling pathway^52^. Similarly, we identified a duplicated and translocated copy of *amhr2* in the 14 kb region. We also identified *gsdf* and *bmpr1bb*, both members of the *TGF-β* signalling pathway, in the SDR of *S. australasicus*, underscoring the recurrent involvement of this pathway in sex determination in *Scomber*, although direct evidence for either gene as the primary sex-determining factor is lacking. In contrast, no TGF-β pathway genes were identified in the SDR of *S. japonicus*, highlighting the flexible, rather than deterministic, nature of the SD gene transitions in this genus.

Our results reveal rapid sex chromosome turnover in *Scomber* genomes. As discussed above, their genomes are characterized by frequent chromosomal rearrangements, particularly on chromosome 24, where both inter- and intraspecific inversions were detected. These features indicate substantial chromosomal instability, which may facilitate further structural rearrangements (e.g., inversions, duplications, translocations) and promote the formation and expansion of nonrecombining regions—key processes that could drive rapid sex chromosome turnover. Future detailed analyses, including epigenomic approaches, such as methylation analyses, will further elucidate the molecular mechanisms underlying chromosomal rearrangements and the expansion of recombination suppression on sex chromosomes.

By integrating long-read and long-range sequencing with comprehensive bioinformatic analyses, we generated high-quality haplotype-resolved genomes for multiple *Scomber* species, enabling fine-scale comparisons of genome structure and evolution both within and between species. Using large-scale population data, we identified extensive chromosomal rearrangements, including megabase-scale inversions, together with rapid sex chromosome turnover, leading to distinct evolutionary trajectories of sex chromosome differentiation and recombination suppression across *Scomber* genomes. While research on fish ecological and evolutionary genomics has focused predominantly on freshwater and coastal species that exhibit pronounced ecological or morphological differentiation, our findings reveal unexpected genomic and evolutionary diversity in highly abundant migratory pelagic fishes that have received relatively little attention. These results highlight the value of such taxa not only as critical resources for sustainable fisheries but also as powerful systems for elucidating the mechanisms and pace of genome evolution in the marine environment.

## Methods

### Ethics statement

All of the animal experiments in this study were performed in compliance with protocols approved by the Institutional Animal Care and Use Committee (IACUC) of Tokyo University of Marine Science and Technology (TUMSAT) and) of the Graduate School of Agricultural and Life Sciences, University of Tokyo. All methods were performed in accordance with the IACUC guidelines and regulations.

### Sample preparation and sequencing for the *de novo* reference genome assembly of a ZW *Scomber japonicus* and an XY *S. australasicus*

Highly contiguous *de novo* genome assemblies of a ZW female *Scomber japonicus* and an XY male *Scomber australasicus* were constructed from PacBio HiFi long-read sequencing and Dovetail Omni-C scaffolding. These samples were offspring of wild-caught parents that were commercially collected in Tateyama (Chiba, Japan). Artificially fertilized eggs were produced in May 2022 at the Tateyama Station of Tokyo University of Marine Science (Chiba, Japan) and transported to the Fisheries Laboratory, University of Tokyo (Shizuoka, Japan) for rearing, as described previously^16^. At 90 days post-fertilization (dpf), a female *S. japonicus* and a male *S. australasicus* were euthanized with an overdose of 2-phenoxyethanol (600 ml/L) (Fujifilm, Osaka, Japan). Whole blood samples were collected from the caudal vein using a K_2_EDTA-treated syringe and sample tubes. Blood samples were immediately frozen in a -80°C freezer until DNA extraction. Moreover their genotypic sex was confirmed using the sex-linked genetic markers described previously^20^ and DNA samples collected as described below.

High-molecular-weight DNA was extracted from an aliquot of blood using a NucleoBond HMW DNA kit (Macherey-Nagel, Düren, Germany) according to the manufacturer’s protocol. DNA quality and quantity were assessed using a TapeStation 4150 system with a Genomic DNA ScreenTape (Agilent Technologies, Tokyo, Japan). The extracted DNA (DIN > 7.0) was sent to Macrogen Corporation Japan (Tokyo, Japan) for HiFi sequencing on a PacBio REVIO system using a single SMRT cell per individual. This process resulted in 76.82 Gb (∼90×) and 51.13 Gb (∼60×) of HiFi reads for female *S. japonicus* and male *S. australasicus*, respectively, with N50 values of 19.48 kb and 19.06 kb, respectively (Supplementary Table S1). Another aliquot of blood was used for Omni-C library construction with a Dovetail Omni-C Kit (Dovetail Genomics, Scotts Valley, CA, USA) using a nonmammalian tissue sample preparation protocol. Briefly, 10 µl of whole blood was treated with disuccinimidyl glutarate (DSG) and formaldehyde for chromatin crosslinking. The crosslinked chromatin was digested with DNase I (1:5 dilution), released from cells in a sodium dodecyl sulfate (SDS) solution, and subjected to end-polishing followed by bridge ligation, proximity ligation, and crosslink reversal. Purified DNA was used for sequencing library construction with NEBNextUltra enzymes and Illumina-compatible adapters. Fragments tagged with biotin during bridge ligation were isolated using streptavidin beads before PCR enrichment. The quality and quantity of the libraries were evaluated using the TapeStation 4150 system with a D1000 ScreenTape kit. The libraries were subsequently sent to GeneBay Inc. (Kanagawa, Japan) for sequencing on a DNBseq-G400RS system (BGI, Shenzhen, China) with a 2×150 bp read length, yielding 121.31 Gb (∼146.4×) and 137.58 Gb (∼165.8×) of reads for *S. japonicus* and *S. australasicus*, respectively (Supplementary Table S11). Methods for the subsequent bioinformatics analysis are explained in the Supplementary information.

### Samples for additional chromosome-level genome assemblies of *S. japonicus* and *S. australasicus*

Additional long-read HiFi sequences were obtained from four individuals of *S. japonicus* and *S. australasicus* (one individual per sex per species) using the PacBio Sequel II system. Individuals of *S. japonicus* were commercially collected in the offshore area of the Bungo Channel (landed Oita, Japan), while *S. australasicus* individuals were collected in the offshore area of the East China Sea (landed Nagasaki, Japan). Phenotypic and genotypic sex was confirmed as outlined above. Whole blood was collected from each individual as described above, and high-molecular-weight genomic DNA was extracted using a NanoBind CBB Big DNA Kit (Circulomics, Baltimore, USA) according to the manufacturer’s instructions. The quality and quantity of each DNA sample were also assessed as described above. Size selection, library preparation, and sequencing were conducted for HiFi sequencing on a PacBio Sequel II at BGI Japan (Tokyo, Japan) for *S. japonicus* and at Macrogen Corporation Japan for *S. australasicus*. Sequencing was performed on a single SMRT cell per individual, resulting in 60.57 Gb (∼72×), 65.76 Gb (∼79×), 45.94 Gb (∼55×) and 51.79 Gb (∼62×) of HiFi sequences with N50 values of 14.3 kb, 14.1 kb, 12.7 kb and 13.3 kb for the ZW, ZZ, XY, and XX individuals, respectively (Table S11). Methods for the subsequent bioinformatics analysis are explained in the Supplementary information. Sample information is summarized in Supplementary Table S14 together with the samples used for the reference genome assembly and Nanopore sequencing for structural variant detection.

### Public datasets used to reassemble *S. japonicus* and *S. scombrus* genomes

Reference genomes of a male *S. japonicus* (fScoJap1) and a *S. scombrus* of unknown sex (fScoSco1.1) have been registered in the NCBI database. Instead of directly using these existing reference genomes, we reassembled the genome from the archived sequencing data associated with these references (SRP470260 for fScoJap1 and ERR12318580/ERR12303949 for fScoSco1.1) using the same strategy as for the *de novo* assembly mentioned above, allowing unbiased comparisons without concerns about differences in assembly methods. The assembly step is explained in the Supplementary information.

### Sex chromosome scanning for *S. japonicus* and *S. australasicus*

A total of 281 (136 males and 145 females) individuals of *S. japonicus* captured from Tatayama, Fukui, and Oita (Japan) and a total of 188 (96 males and 92 females) individuals of *S. australasicus* captured from Tateyama and Nagasaki (Japan) were used (Supplementary Table S5) for the identification and characterization of sex chromosomes. Methods for sequencing and the subsequent bioinformatics analysis, including genotyping, the detection of sex-associated loci and population genetics analyses, are explained in the Supplementary information.

### Location of the sex-determining region of *S. scombrus*

RAD-seq data from 158 wild males and 143 wild females were obtained from the NCBI Sequence Read Archive (SRP491935) to identify the sex-linked regions in the *S. scombrus* genome. Methods for the subsequent bioinformatics analysis, including SNP genotyping and the detection of sex-associated loci, are explained in the Supplementary information. Sex-diagnostic codominant markers, which amplify both the male-specific region and its paralogous autosomal region, were designed utilizing the BLASTn results (Supplementary Table S12). Genomic DNA was extracted from 16 females and 14 males captured offshore of the Atlantic coast of Portugal using the DNeasy Blood and Tissue Kit (Qiagen) according to the manufacturer’s protocol. PCRs were performed in a total reaction volume of 25 μl containing 2 μl (10 μM) of each primer, 12.5 µl of Hot-Start GoTaq DNA Polymerase Master Mix (Promega), 6.5 µl of nuclease-free water, and 2 µl (10 ng/µl) of genomic DNA. The following thermal conditions were used: initial denaturation at 94°C for 5 min; 30 cycles at 94°C for 60 s, 58°C for 30 s and 72°C for 1 min; and a final one-cycle elongation step at 72°C for 5 min. Amplicon sizes were visualized by electrophoresis using a D1000 ScreenTape kit on a 4150 TapeStation instrument (Agilent). Methods for the identification of heterogametic SD regions are described in the Supplementary information.

## Data Availability

The haplotype-resolved assemblies of ZW female *S. japonicus* and XY male *S. australasicus* have been deposited in the DNA Data Bank of Japan (DDBJ). The assemblies of *S. japonicus* are available under the umbrella BioProject **********, with ********** for the primary haplotype (sequence accession numbers ********–********) and ********** for the alternate haplotype (sequence accession numbers ********–********). The assemblies of *S. australasicus* are available under the umbrella BioProject **********, with ********** for the primary haplotype (sequence accession numbers ********– ********) and ********** for the alternate haplotype (sequence accession numbers ********– ********).

## Code Availability

All the data were analysed using publicly available software and packages in this study. All the software and packages used in this study are described in the Supplementary information file.

## Acknowledgements

We are grateful to Dr. Motoshige Yasuike for sharing the protocol for Omni-C library construction. We also thank the members of Tokyo University of Marine Science and Technology. This study was supported by the Japan Science and Technology Agency (JST) Mirai Program (JPMJMI18CH and JPMJMI21C1) and the Japan Society for the Promotion of Science (JSPS) Grant-in-Aid for Scientific Research (B) (21H02279).

## Competing interests

The authors declare that they have no competing interests.

## Supplementary Information

- The Supplementary information file includes the Supplementary figures and methods.
- Supplementary Tables S1–S14 (.xlsx)

## Notes

### Competing Interest Statement

The authors have declared no competing interest.

