## Supplementary figures and methods for "Rapid chromosomal rearrangements and sex chromosome turnover underlie the evolution of parapatric Pacific *Scomber* mackerels"


**Author Information**

Ahammad Kabir[1], Ryosuke Yazawa[2], Dana Marielba Silva[3], Jorge M.O. Fernandes[3], Masaomi Hamasaki[4], Sota Yoshikawa[4], Hiroaki Suetake[5], Kiyoshi Kikuchi[1], and Sho Hosoya[1,*]

*Corresponding author

**Affiliations**

1 Fisheries Laboratory, Graduate School of Agricultural and Life Sciences, University of Tokyo, Shizuoka, Japan

2 Department of Marine Biosciences, Tokyo University of Marine Science and Technology, Tokyo, Japan

3 Institut de Ciències del Mar, Spanish Research Council, Barcelona, Spain

4 Nagasaki Prefectural Institute of Fisheries, Nagasaki, Japan

5 Faculty of Marine Science and Technology, Fukui Prefectural University, Fukui, Japan

#### Contents

**Supplementary Figs. 1–17**

**Supplementary Methods**

#### Supplementary figures

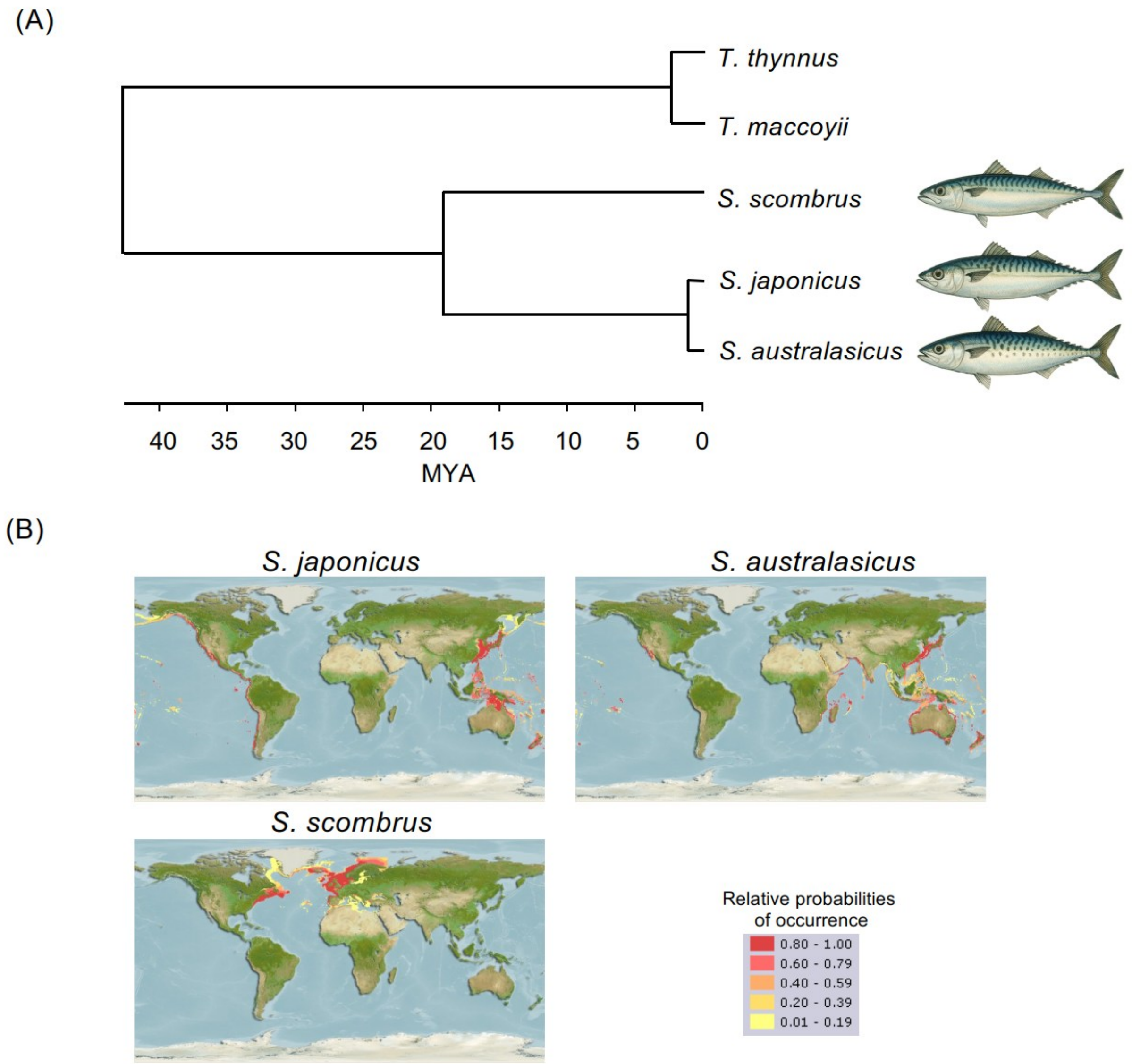


**Supplementary Fig. 1. (A)** Phylogenetic relationships of *Scomber* species—*S. japonicus* and *S. australasicus*, and *S. scombrus*—with two outgroup species, *Thunnus thynnus* and *T. maccoyii*, obtained from TimeTree 5 (https://timetree.org/)[1]. Divergence times were estimated based on fossil records and mitochondrial divergence. Images of *Scomber* species were generated using ChatGPT. **(B)** Global distribution ranges of the four *Scomber* species based on AquaMaps (accessed May 1, 2025) [2]. The distribution of *S. japonicus* reported previously[2,3] has been revised, and Atlantic specimens previously identified as *S. japonicus* were reclassified as *S. colias*[4].

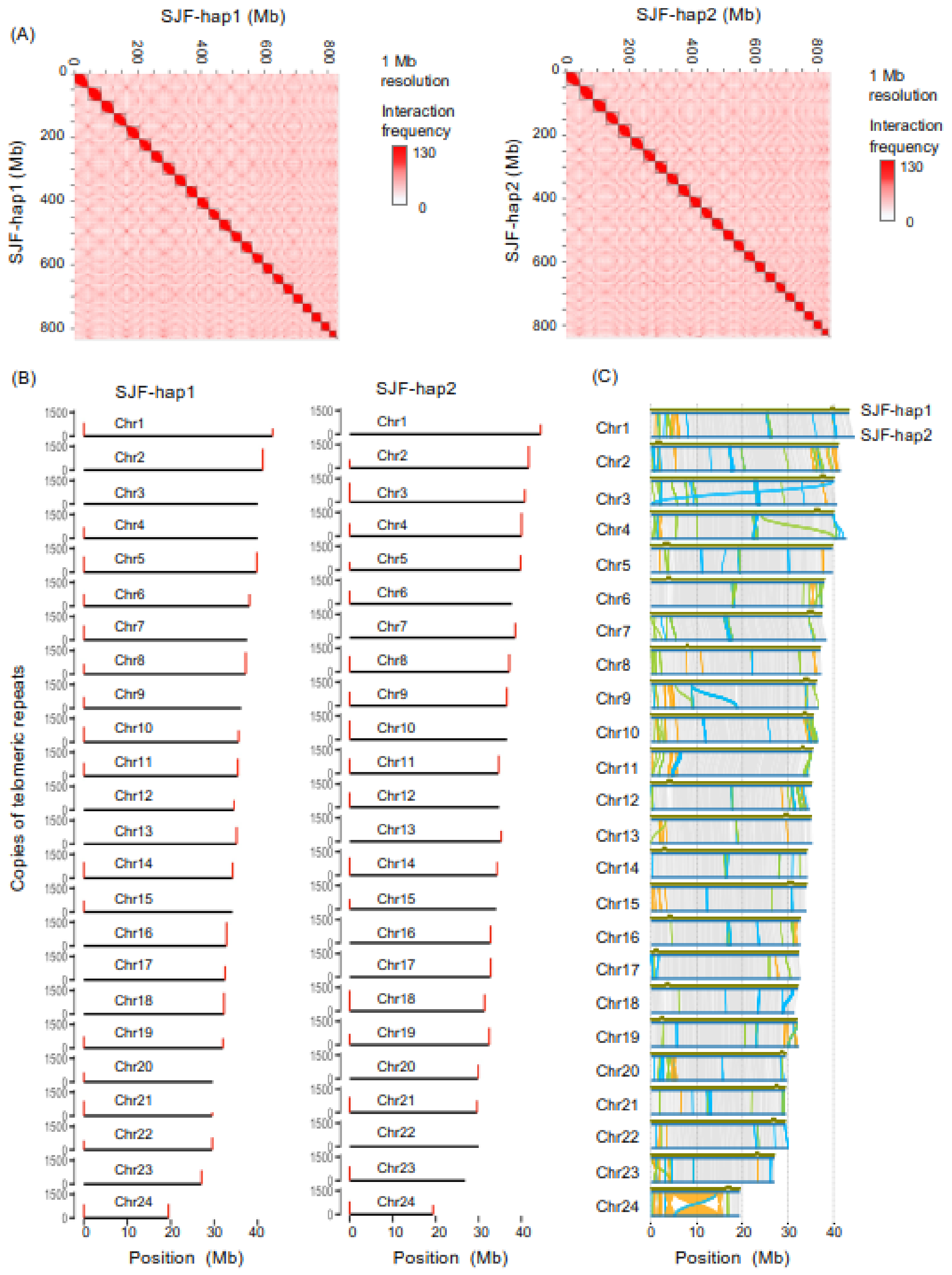


**Supplementary Fig. 2.** Haplotype-resolved genome assemblies of a ZW female *Scomber japonicus* (SJF). **(A)** Omni-C contact maps (1-Mb resolution) for haplotype 1 (SJF-hap1) and haplotype 2 (SJF-hap2) across all 24 chromosomes. **(B)** Copy number of telomeric repeats for each haploid chromosome (left panel: SJF-hap1; right panel: SJF-hap2) along with the telomere localization. **(C)** Synteny between the two haplotypes of the SJF genome. Syntenic, inverted, translocated, and duplicated regions are shown in grey, yellow, green, and sky blue, respectively. Centromere positions on SJF-hap1 are indicated in grey along the green horizontal line at the top of each synteny plot.

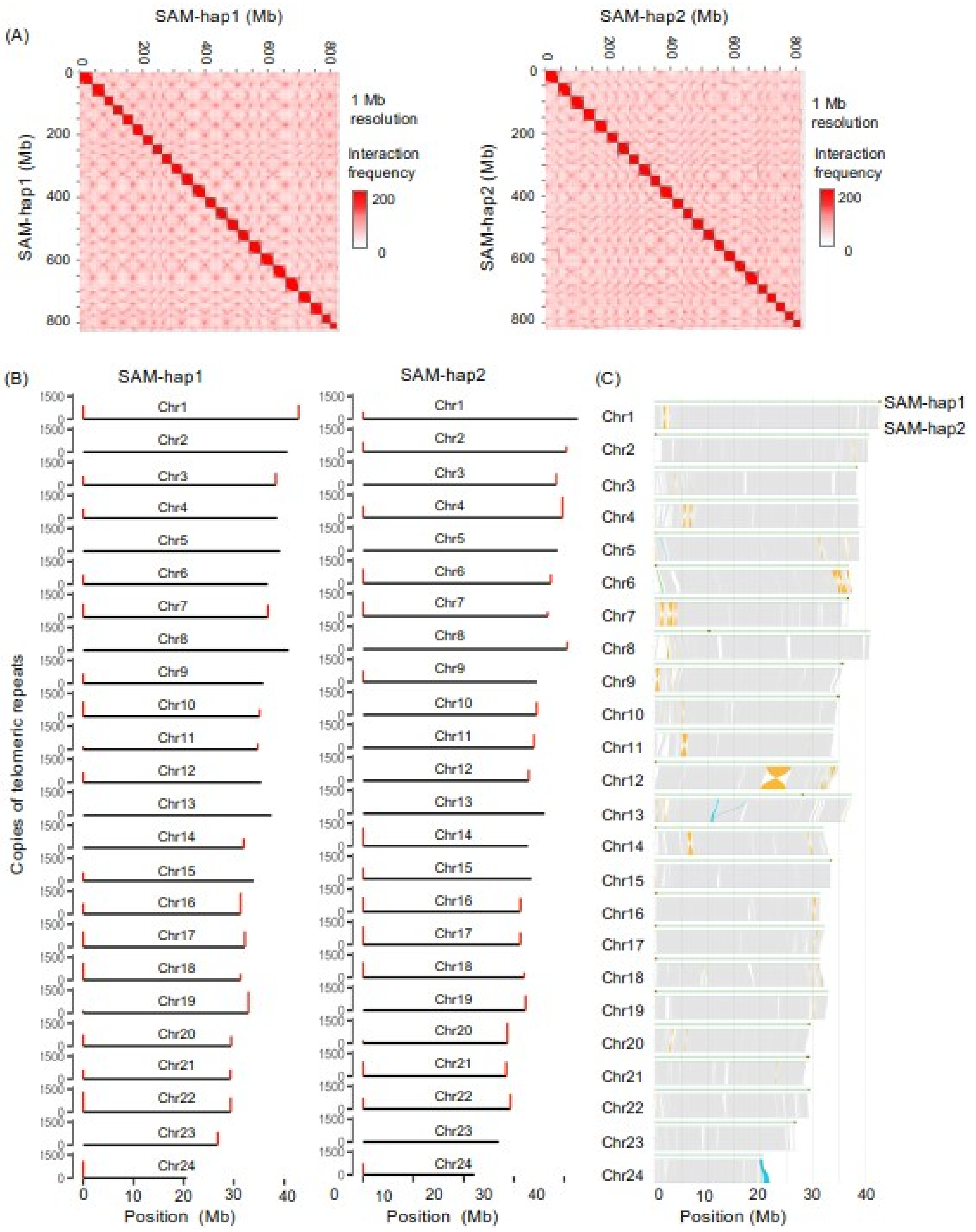


**Supplementary Fig. 3.** Haplotype-resolved genome assemblies of an XY male *Scomber australasicus* (SAM). **(A)** Omni-C contact maps (1-Mb resolution) for haplotype 1 (SAM-hap1) and haplotype 2 (SAM-hap2) across all 24 chromosomes. **(B)** Copy number of telomeric repeats for each haploid chromosome (left panel: SAM-hap1; right panel: SAM-hap2) along with the telomere localization. **(C)** Synteny between the two haplotypes of the SAM genome. Syntenic, inverted, translocated, and duplicated regions are shown in grey, yellow, green, and sky blue, respectively. Centromere positions on SAM-hap1 are indicated in grey along the green horizontal line at the top of each synteny plot.

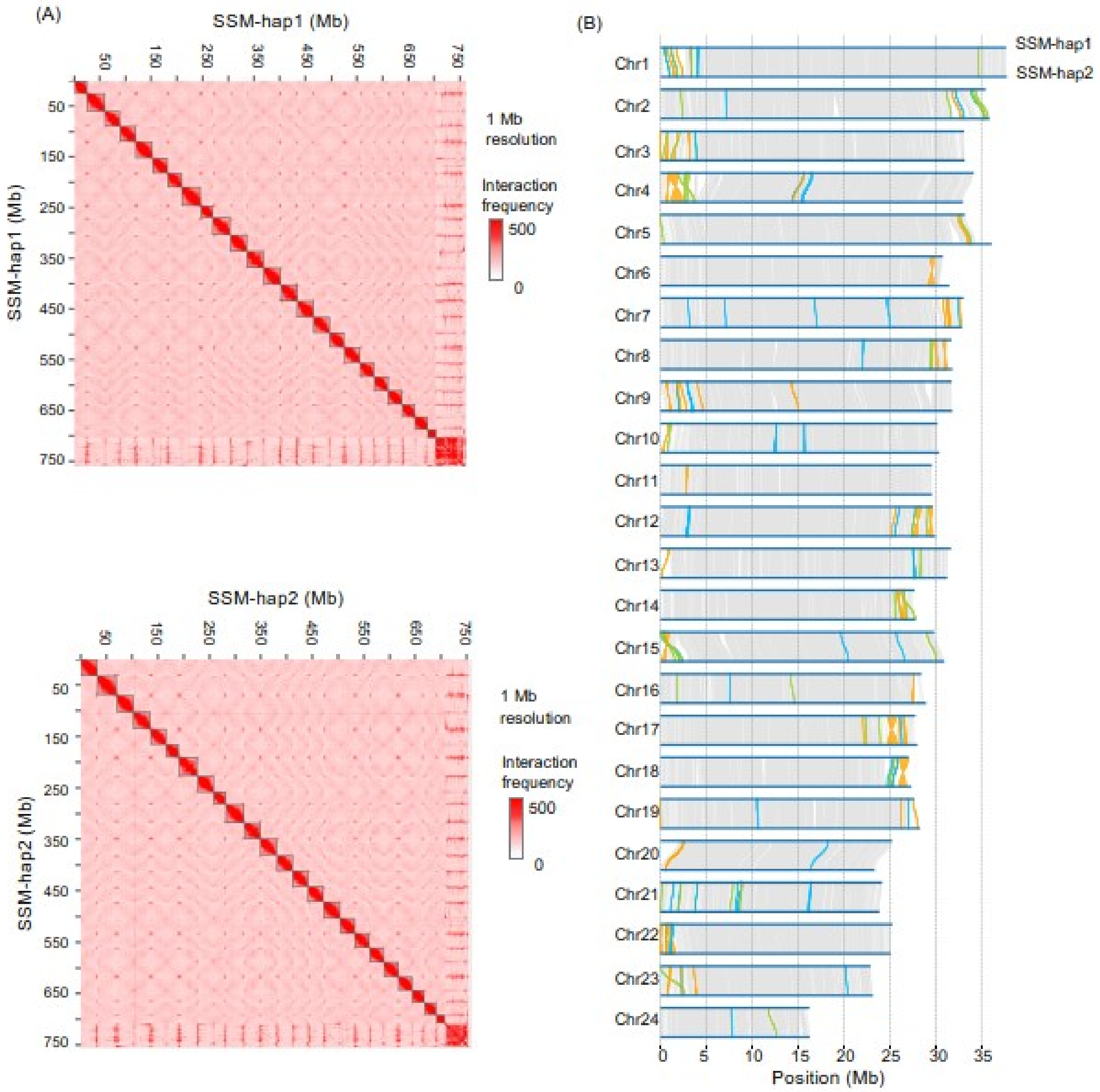


**Supplementary Fig. 4.** Haplotype-resolved genome assemblies of an XY male *Scomber scombrus* (SSM). **(A)** Hi-C contact maps (1-Mb resolution) for haplotype 1 (SSM-hap1) and haplotype 2 (SSM-hap2) across all 24 chromosomes. **(B)** Synteny between the two haplotypes of the SSM genome. Syntenic, inverted, translocated, and duplicated regions are shown in grey, yellow, green, and sky blue, respectively.

Supplementary Fig. 5

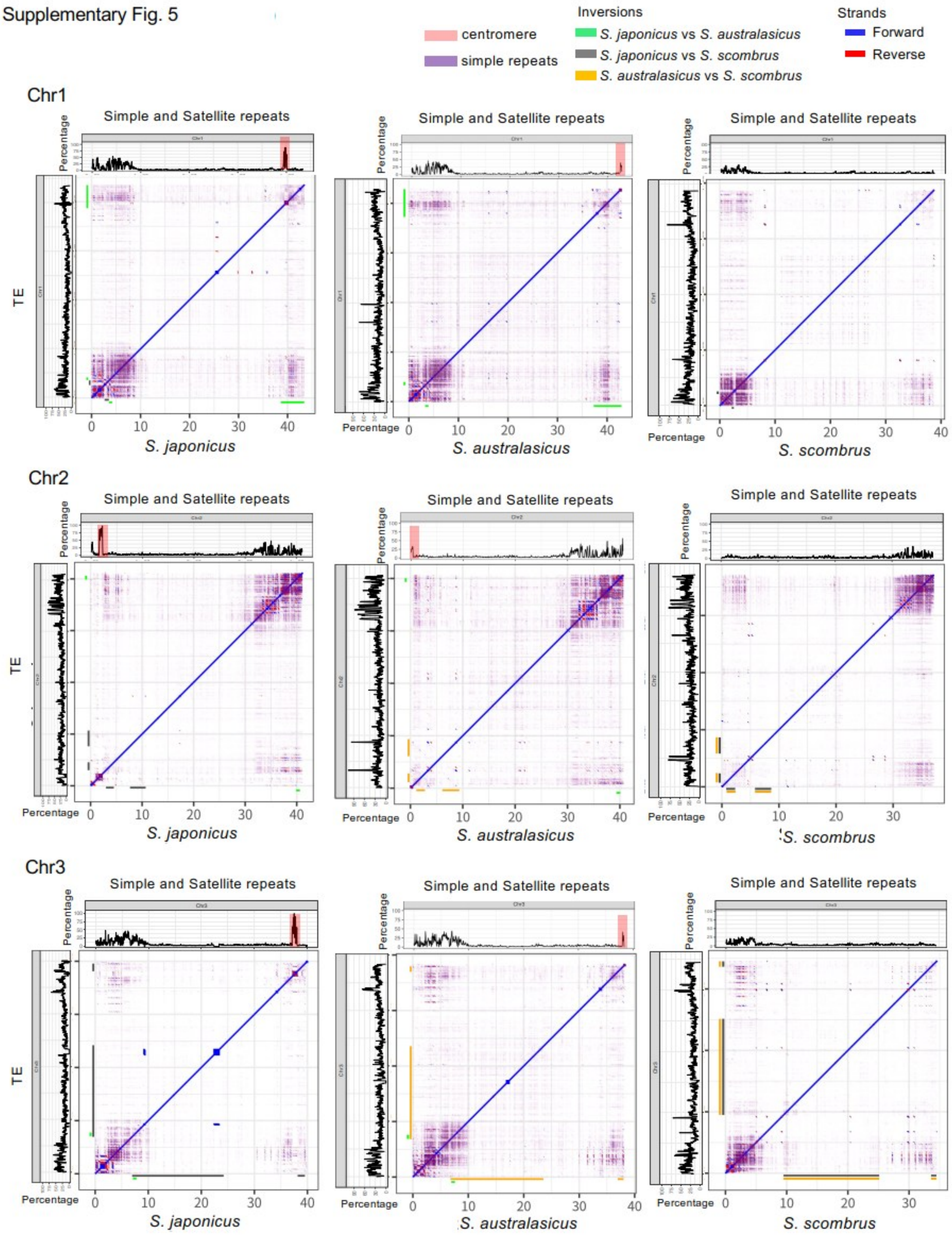

Supplementary Fig. 5 (continued)

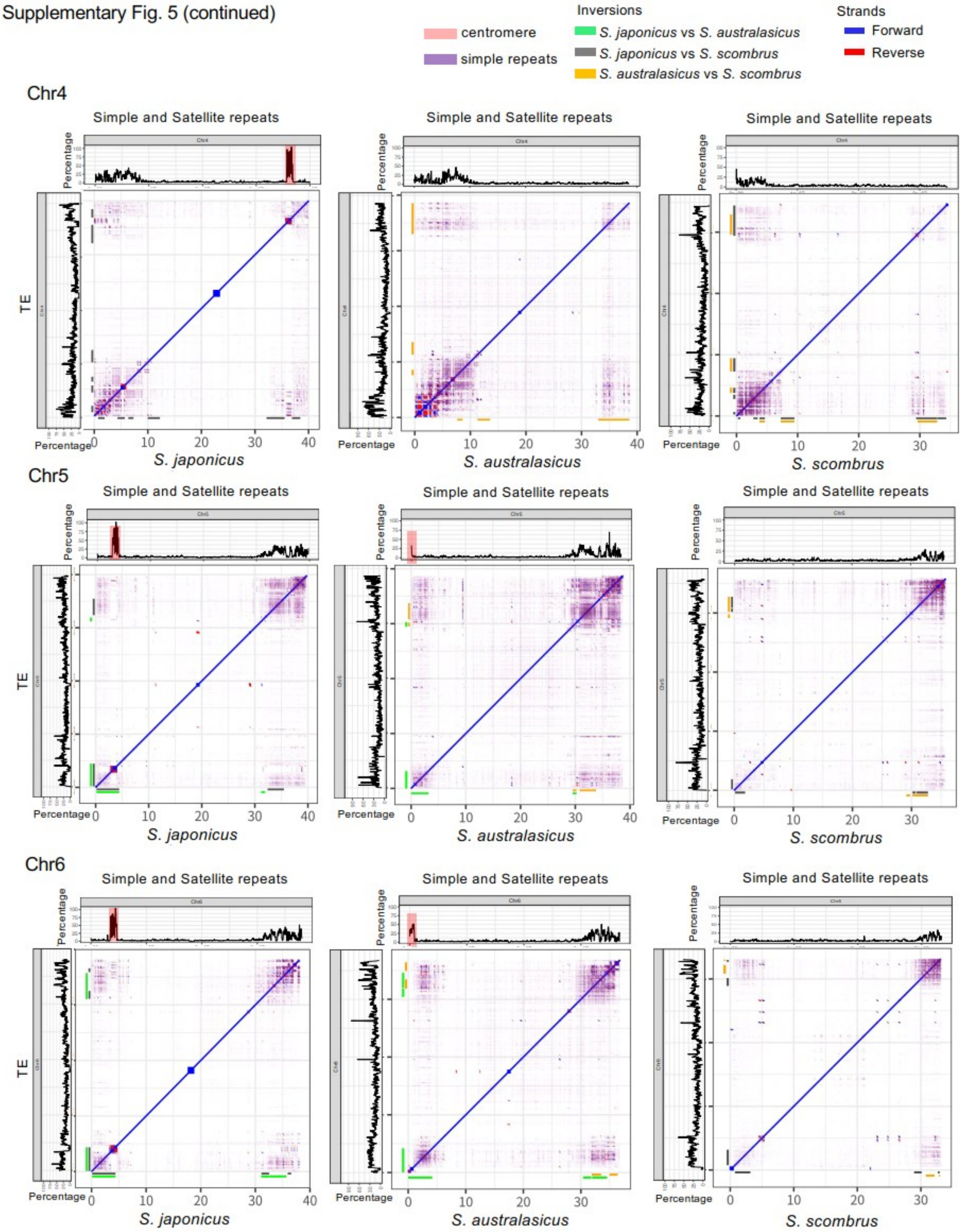

Supplementary Fig. 5 (continued)

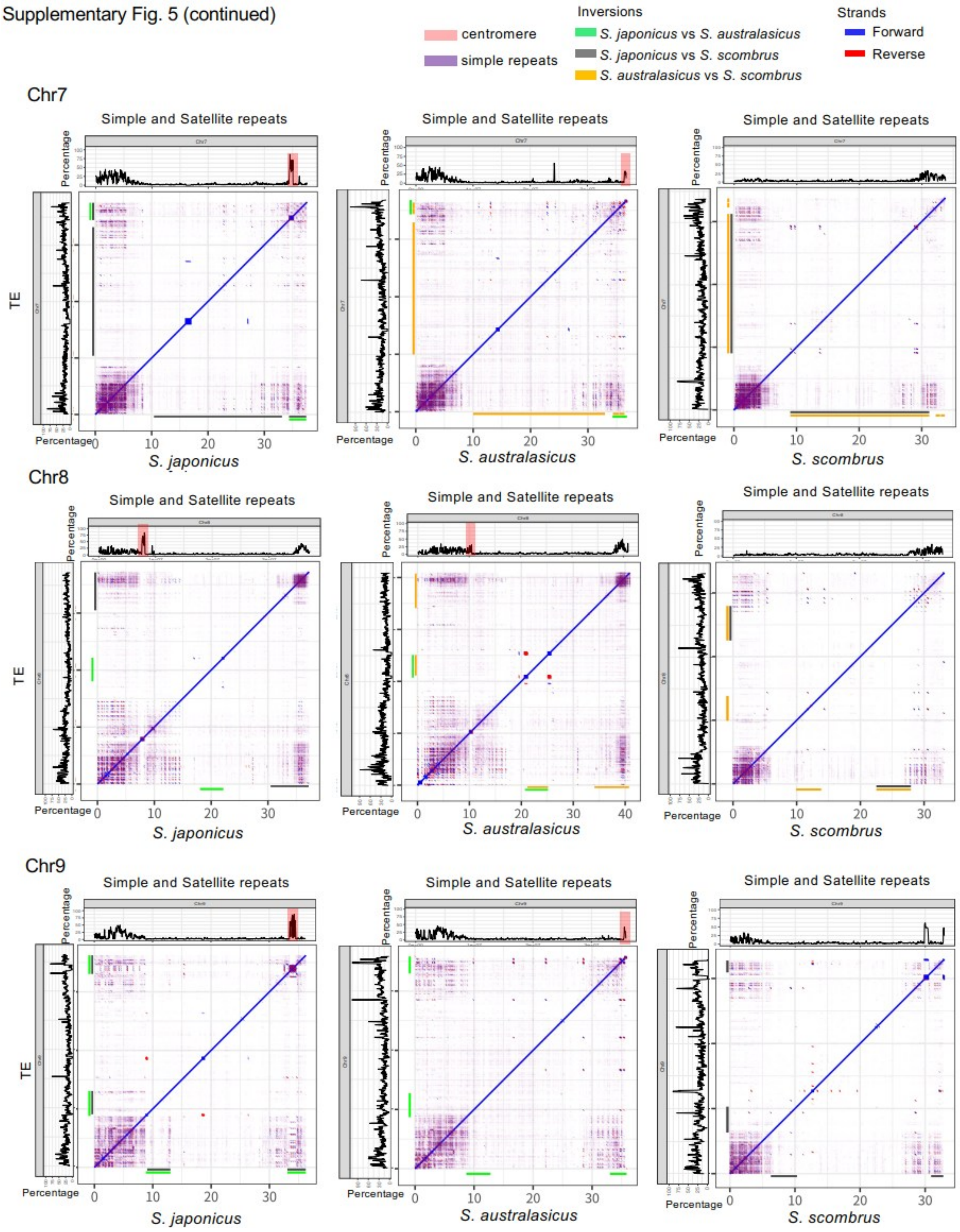

Supplementary Fig. 5 (continued)

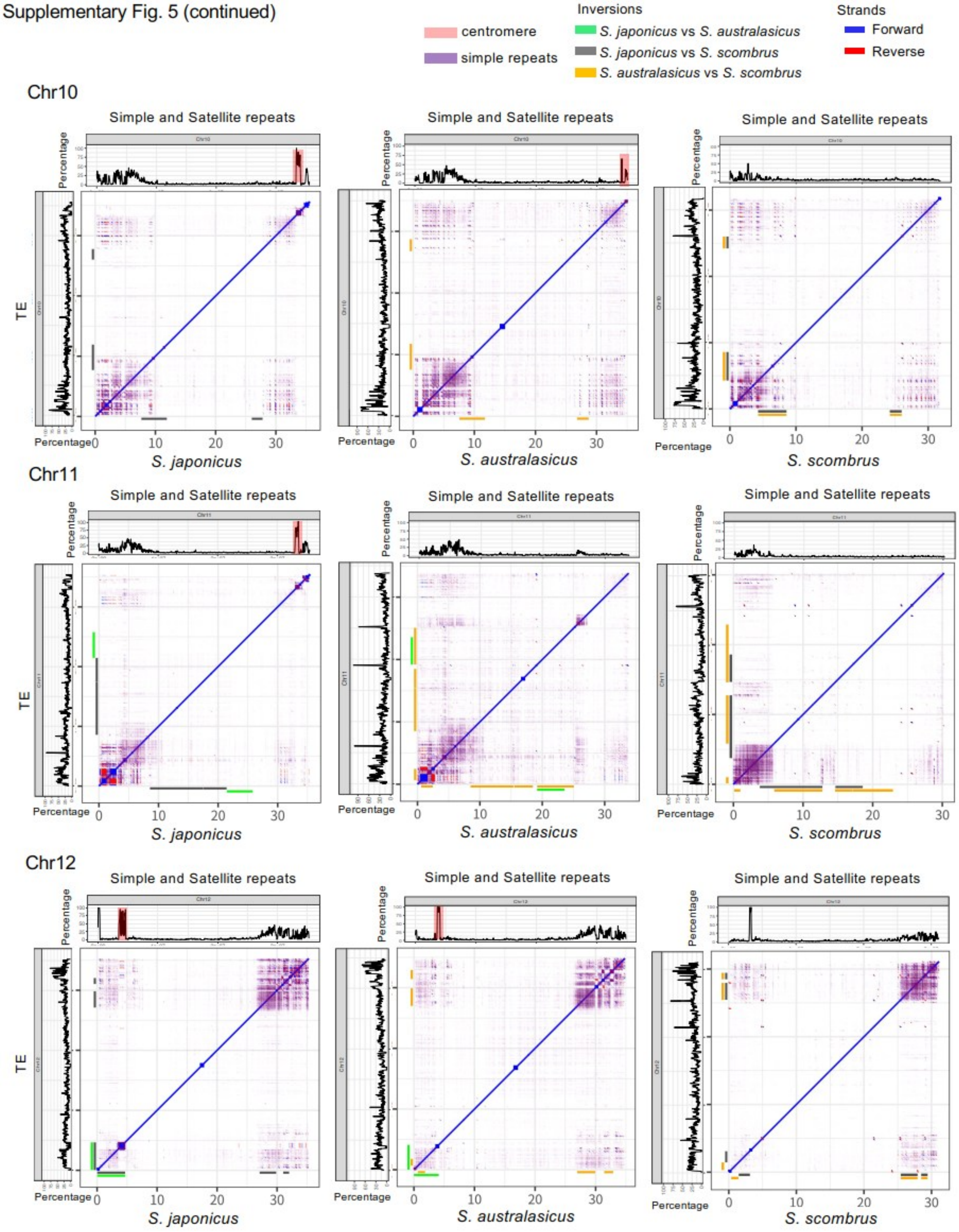

Supplementary Fig. 5 (continued)

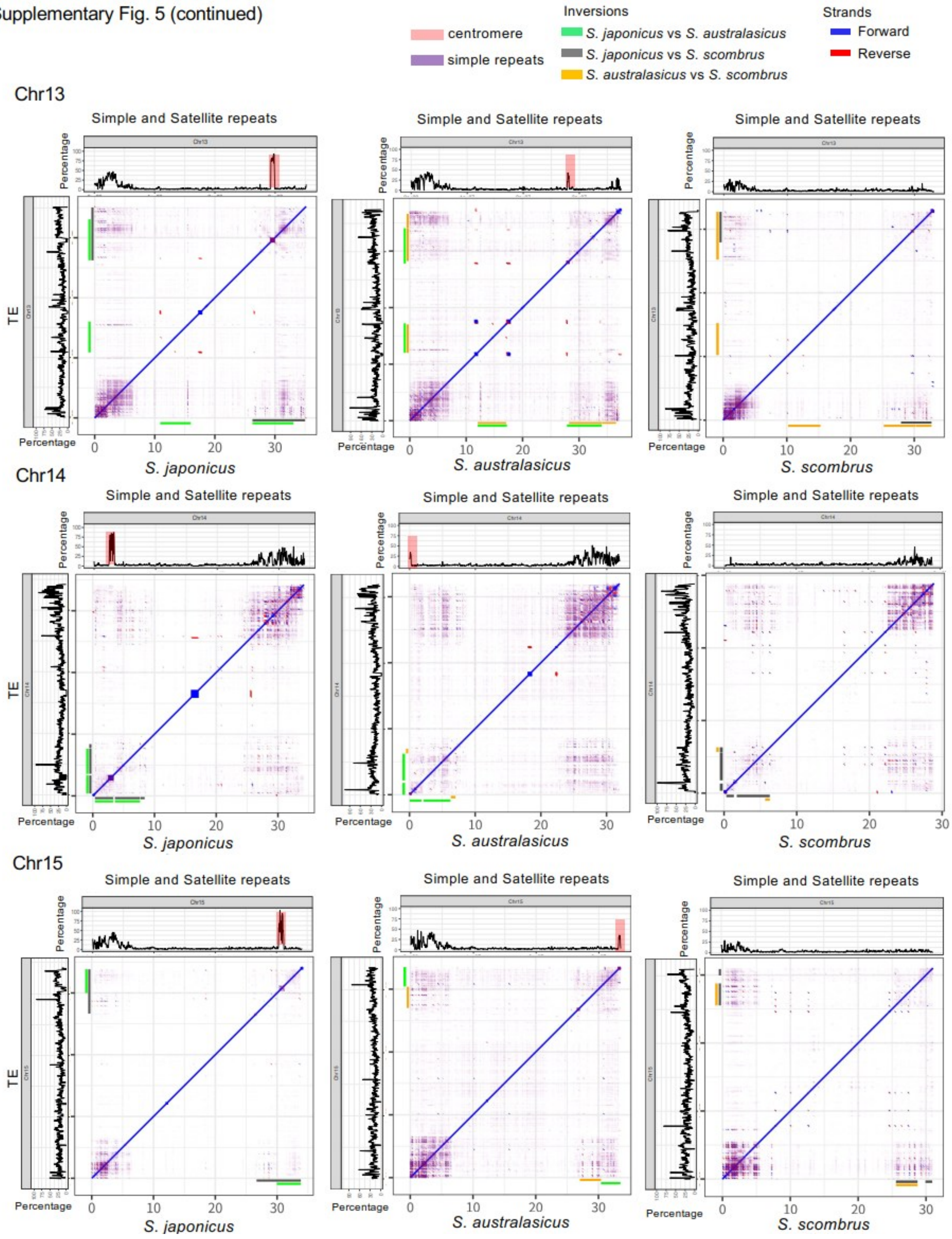

Supplementary Fig. 5 (continued)

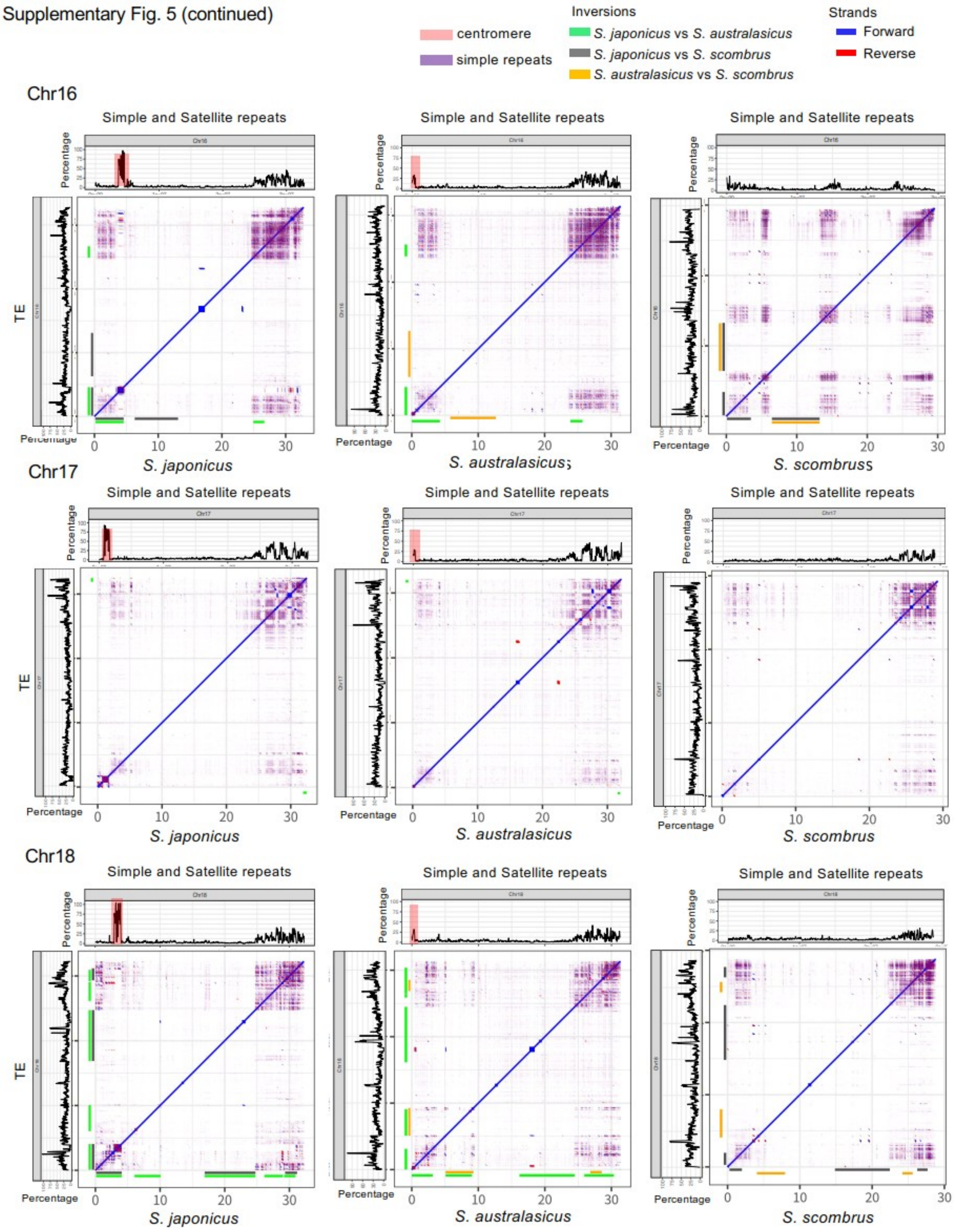

Supplementary Fig. 5 (continued)

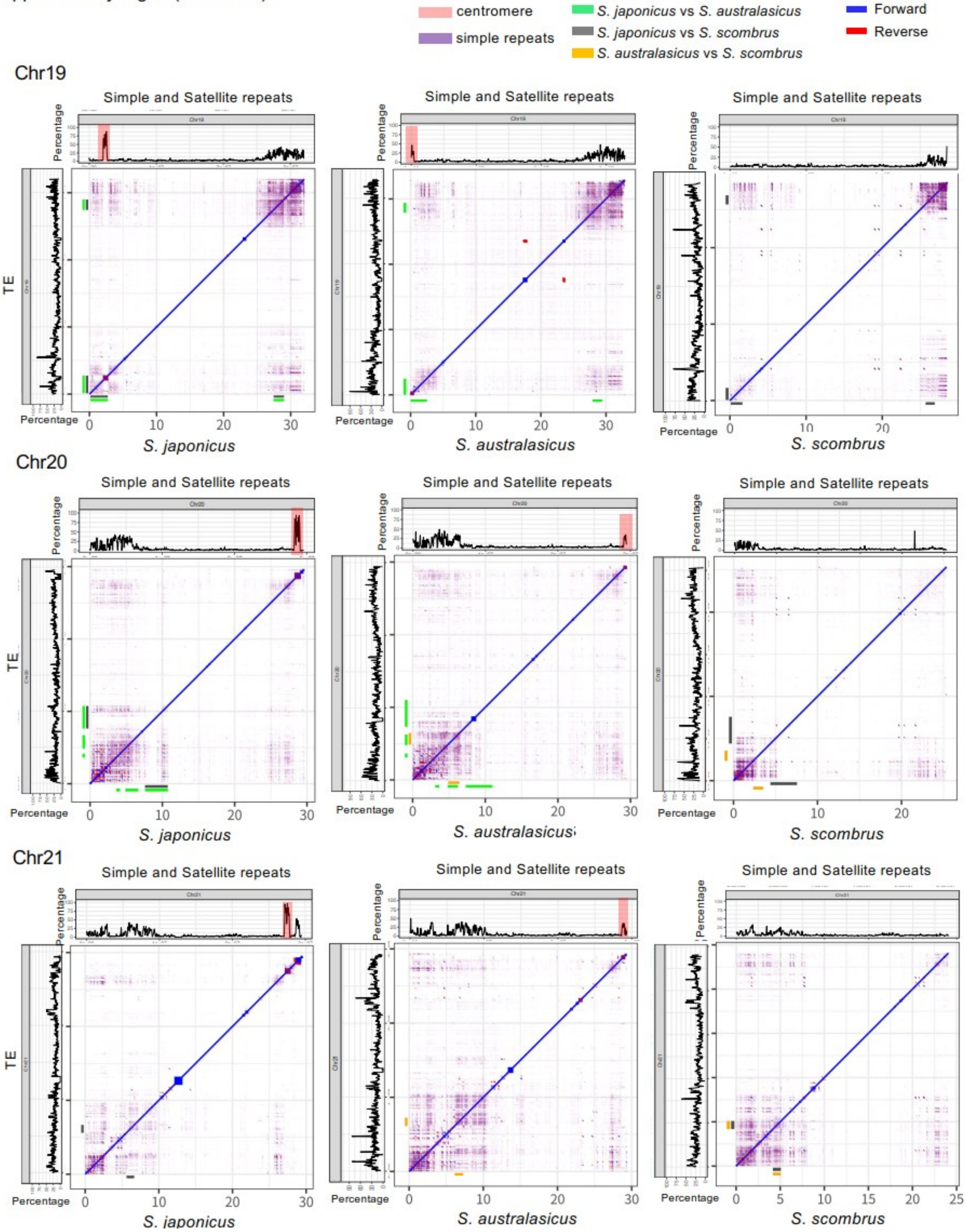

Supplementary Fig. 5 (continued)

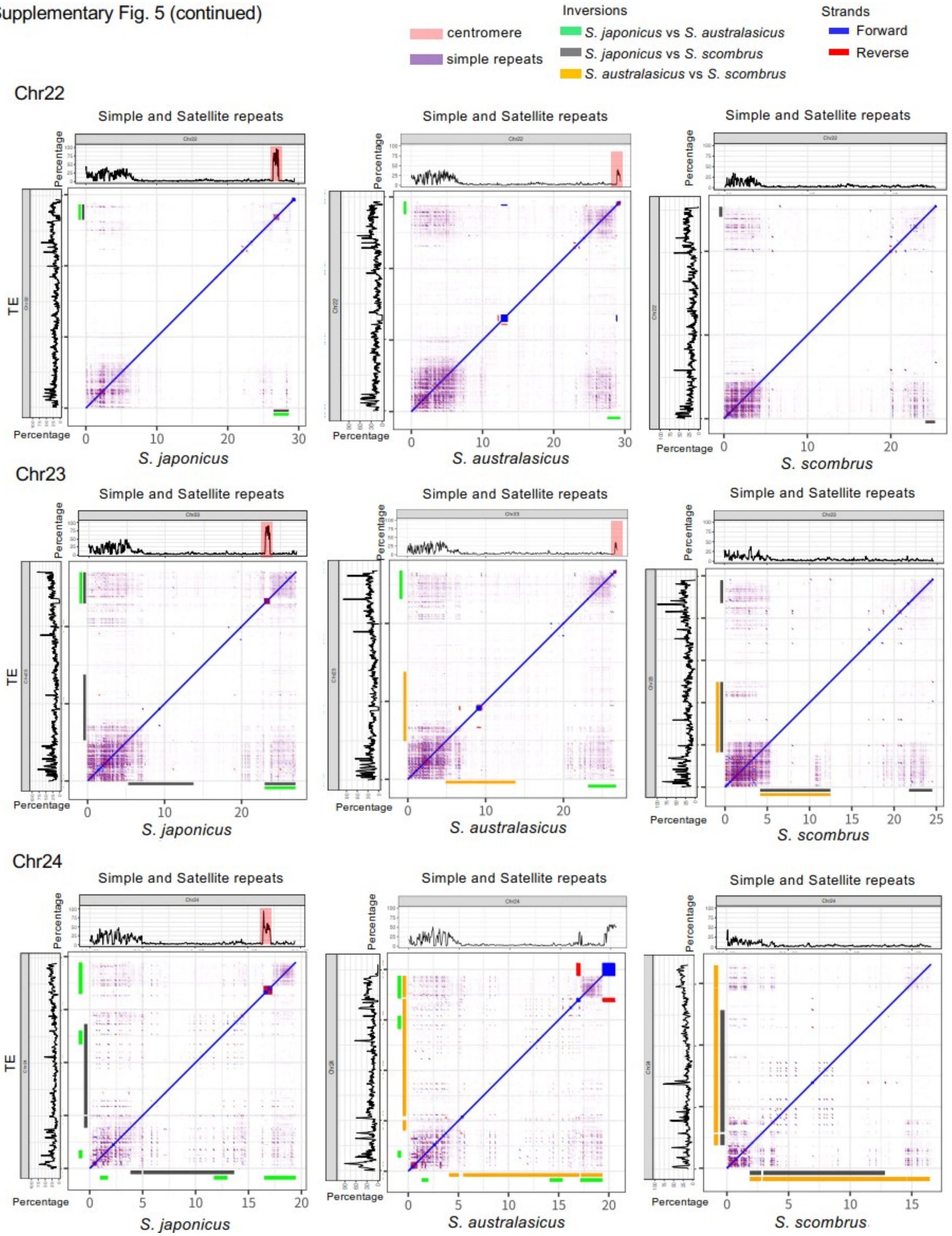

**Supplementary Fig. 5.** Self-aligned dot plots of chromosomes harbouring inversions in *Scomber japonicus* (left panel), *S. australasicus* (middle panel), and *S. scombrus* (right panel). The proportion of genomic sequences masked as simple or satellite repeats in 50-kb windows is shown above each dot plot as a line plot, with centromere positions highlighted in pink. On the left side of each dot plot, the proportion of genomic sequences masked as transposable elements (TEs) in 50-kb windows is shown as a line plot.

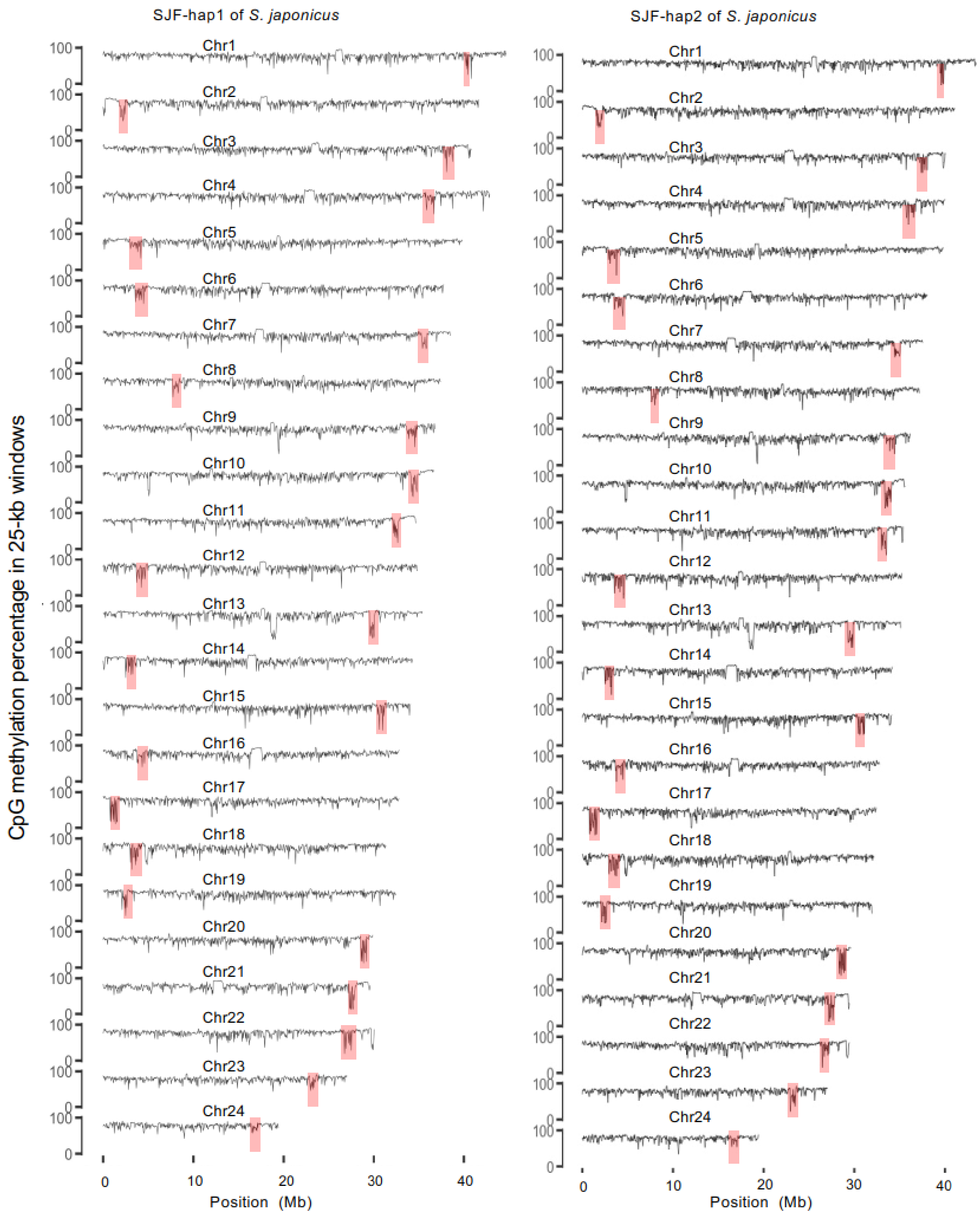


**Supplementary Fig. 6.** CpG methylation levels (%) across the 24 chromosomes of *Scomber japonicus* (SJF-hap1 and SJF-hap2) were calculated in 25-kb windows. Hypomethylated regions (red boxes) indicate the predicted locations of active centromeres on each chromosome.

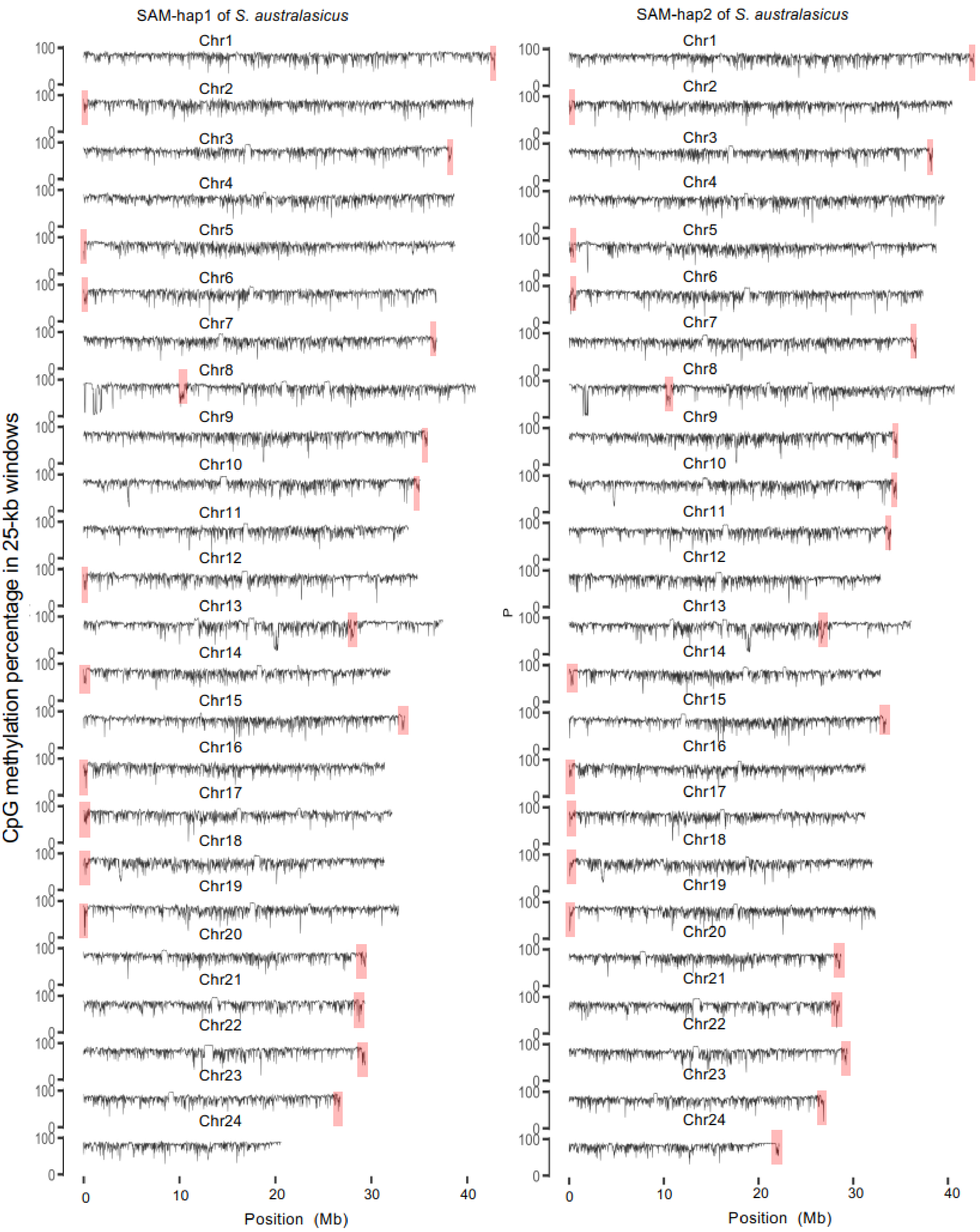


**Supplementary Fig. 7.** CpG methylation levels (%) across the 24 chromosomes of *Scomber australasicus* (SAM-hap1 and SAM-hap2) were calculated in 25-kb windows. Hypomethylated regions (red boxes) indicate the predicted locations of active centromeres on each chromosome.

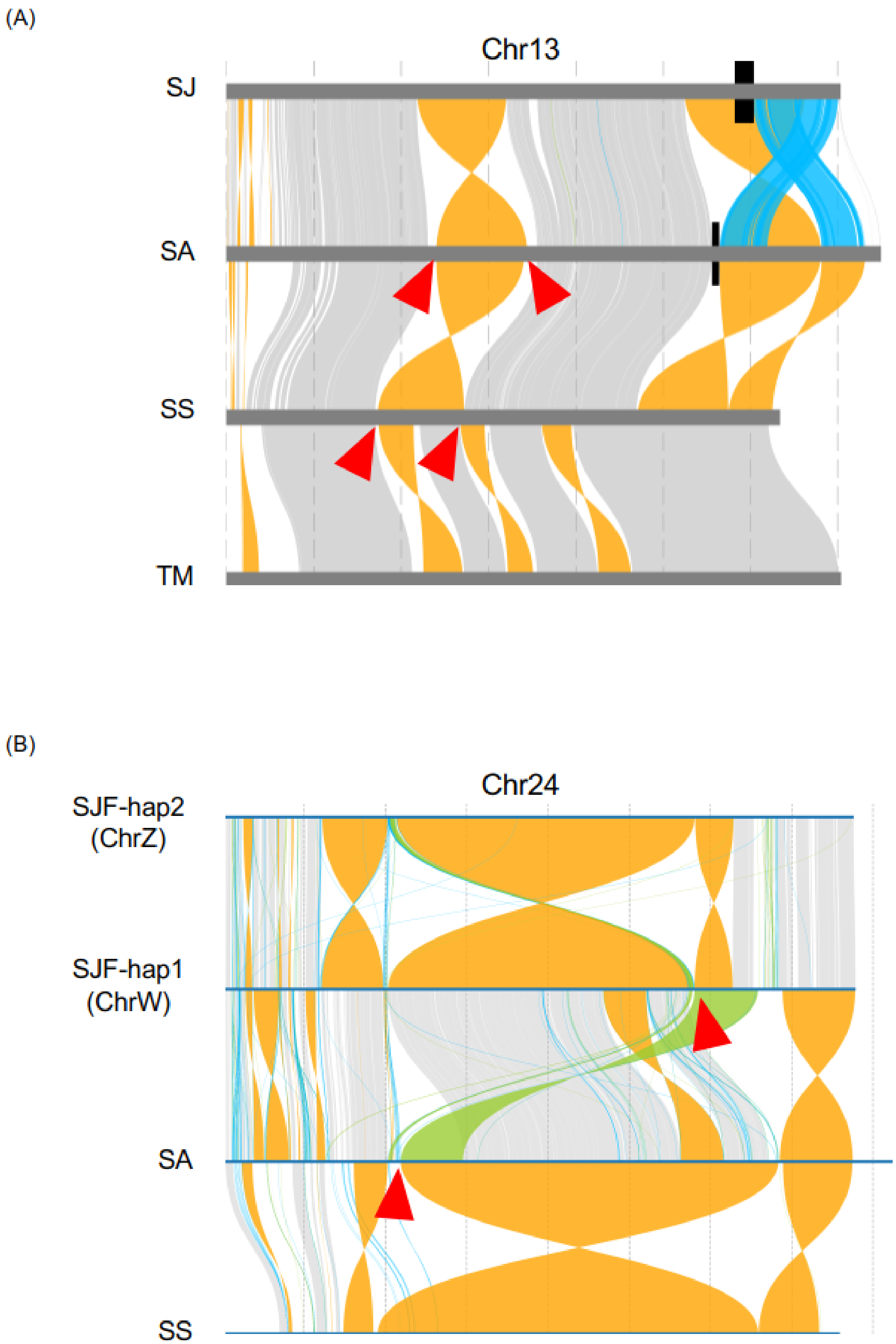


**Supplementary Fig. S8.** Recurrent use of inversion breakpoints. (A) An example on Chr13 is shown; the two breakpoints of an inversion that occurred between *Scomber japonicus* (SJ) and *S. australasicus* (SA), indicated with red sagittal arrowheads, reused one end of the two inversions occurred between *Thunnus maccoyii* (TM) and the common ancestor of *S. scombrus* (SS), SA, and SJ. (B) An example on Chr24 is shown; one end of a breakpoint of inversion that occurred between SJ and SA, indicated with a red sagittal arrowhead, reused the breakpoint of translocation that occurred between SS and the common ancestor of SA and SJ.

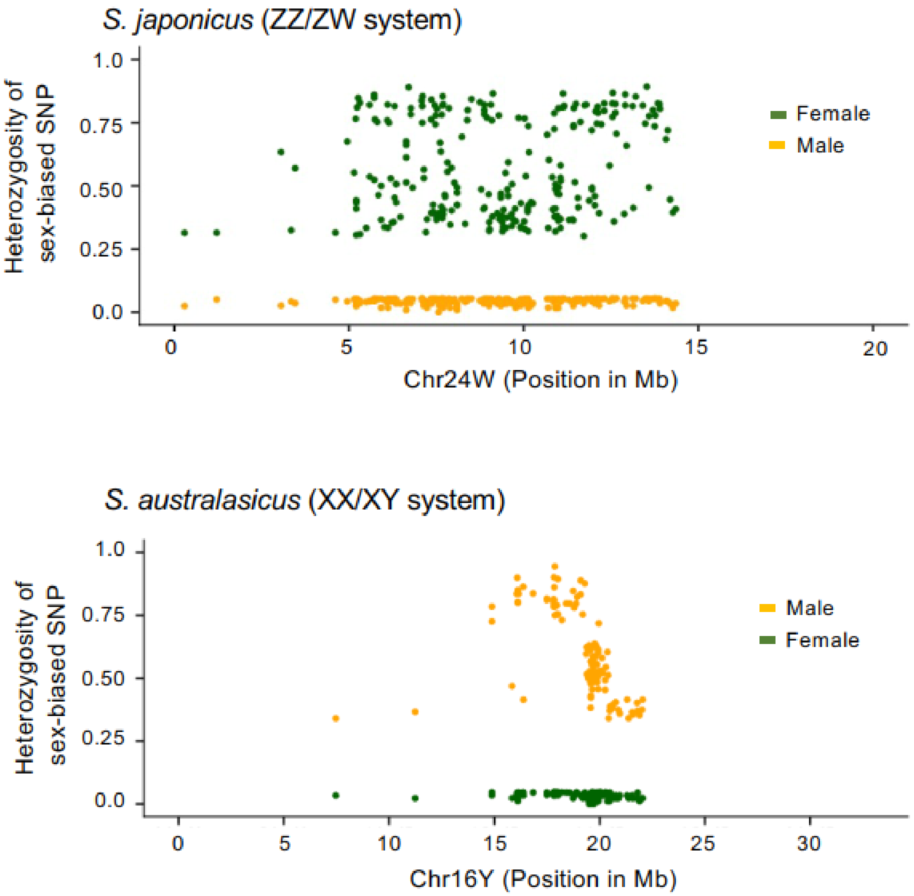


**Supplementary Fig. 9.** Heterozygosity of the sex-biased SNPs for *S. japonicus* (top panel) and *S. australasicus* (bottom panel).

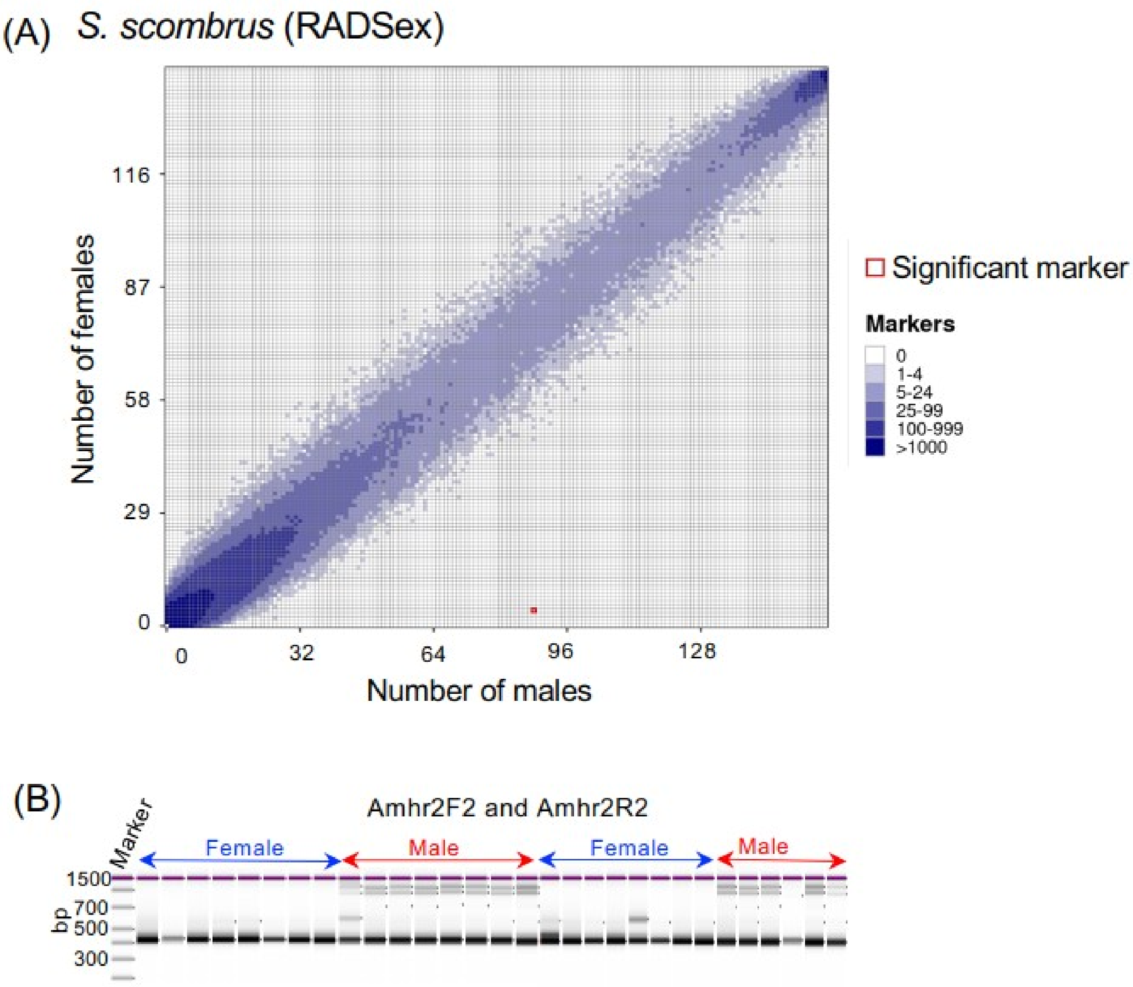


**Supplementary Fig. 10.** Identification of sex-determining (SD) regions in *Scomber scombrus*. **(A)** Identification of the SD locus using RAD-Seq from 158 male and 143 female individuals to identify the SD region in *S. scombrus* with the RADSex approach. The tile plot depicts the distribution of RADSex markers between phenotypic males (x-axis) and phenotypic females (y-axis), with the tile colour intensity corresponding to the number of markers detected in the respective sexes. The tile exhibiting a significant association with phenotypic sex (chi-square test, $p < 0.05$ after the Bonferroni correction) is outlined in red. A single marker was identified in 86 of the 158 males and in 4 of the 143 females, consistent with the XX/XY system in *S. scombrus*. **(B)** Genotyping by PCR using designated primer pairs to distinguish male and female *S. scombrus*.

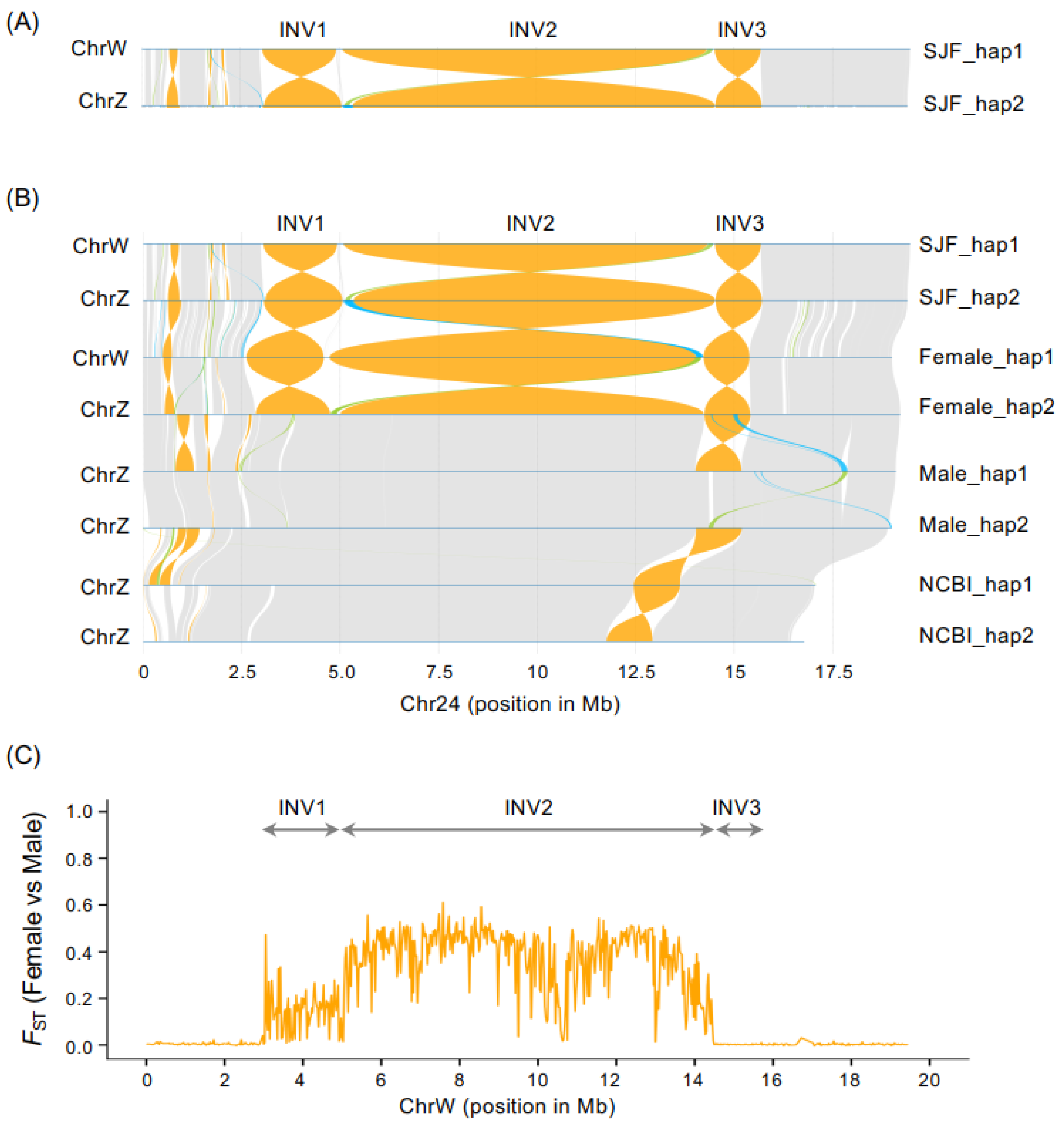


**Supplementary Fig. 11**. **(A)** The synteny plot between ChrW and ChrZ generated from the haplotype-resolved assembly of thefemale individual of *S. japonicus* (SJF). **(B)** The synteny plot of Chr24 using multiple ChrW and ChrZ sequences generated from haplotype-resolved assembly from one additional female and two unrelated male individuals of *S. japonicus*. **(C)** Sequence diversity ($F_{ST}$) between 146 female and 135 male individuals on ChrW.

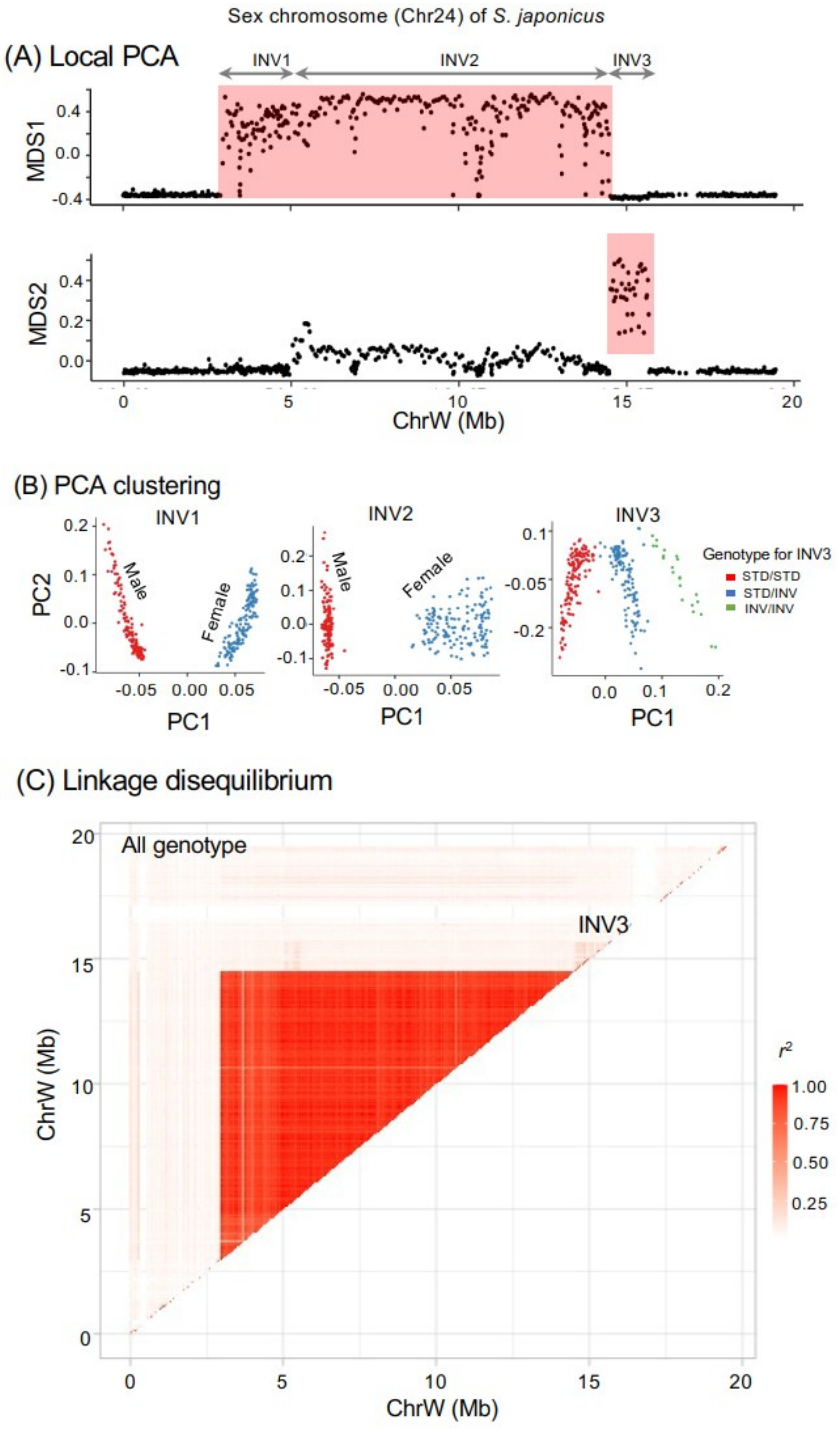


**Supplementary Fig. 12.** Detection of structural variation in Chr24 of *S. japonicus*. **(A)** Local PCA with 100-kb window structural variation consistent with the three inversions detected in the synteny analysis. Distances between local PCA maps are shown on the MDS1 and MDS2 axes, with outlier windows shaded in red. **(B)** Clustering of samples by PCA for INV1, INV2 and INV3, assigned using *k*-means clustering. **(C)** LD across Chr24, shown as maximum $r^2$ values for paired windows. Maximum $r^2$ values are shown for all genotypes from the PCA clustering of the INV3 region. Scales for $r^2$ values are provided (left side of the plot).

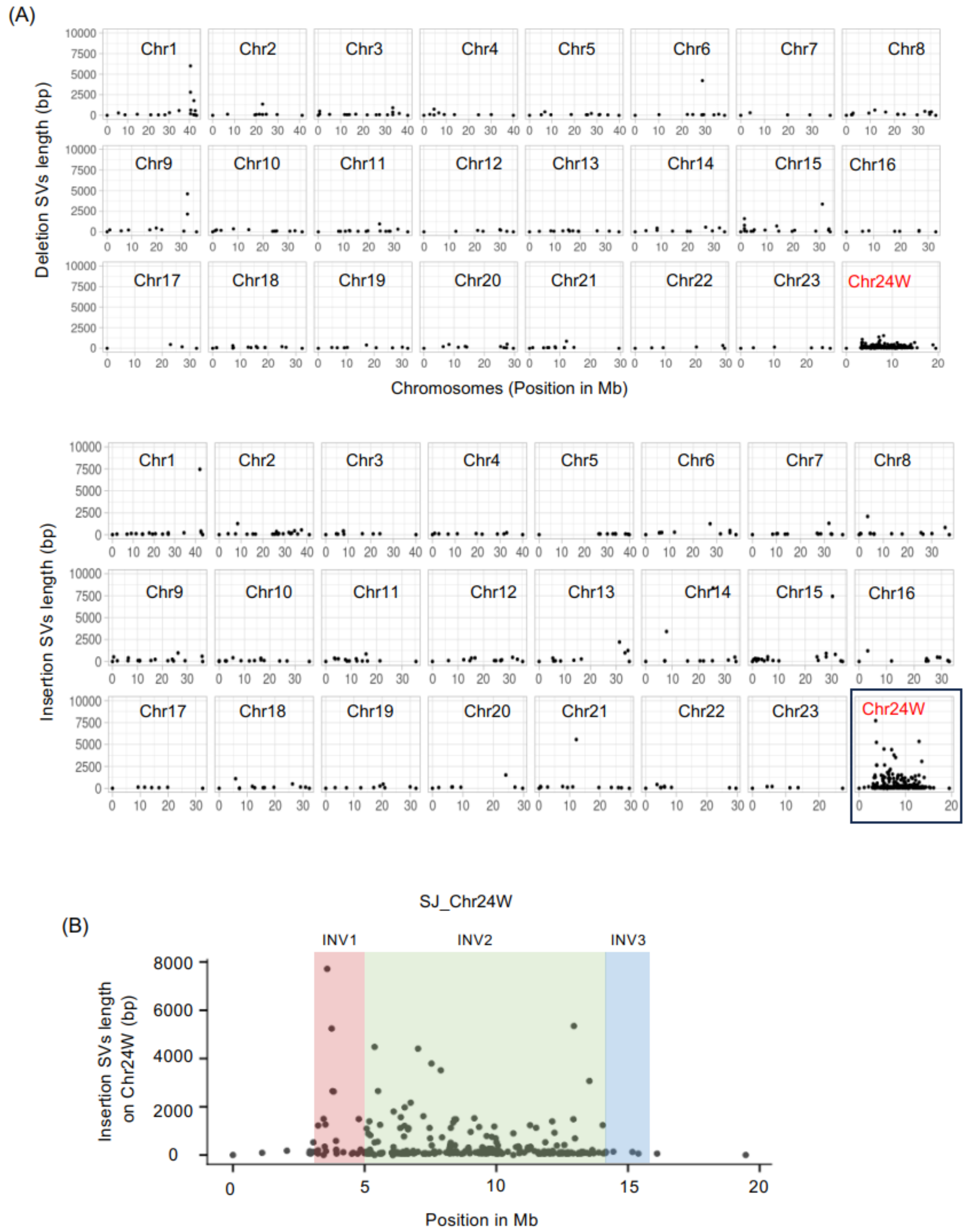


**Supplementary Fig. 13.** Insertion and deletion structural variants (SVs) are enriched on Chr24W. Structural variant calling using read-based alignment. For read-based structural variant calling, we used the HiFi log-reads from two female individuals and one male individual and the Oxford Nanopore long-reads from one male and one female individual. We mapped those reads onto the SJF-hap1 reference genome. We identified female-biased insertion and deletion SVs (> 50 bp) that were heterozygous for females but homozygous for males. (**A**) The identified deletion SVs are shown for 24 chromosomes in the top plot, and the insertion SVs are shown in the bottom plot. The lengths of the SVs are shown on the y-axis, and the chromosomes are shown on the x-axis. (**B**) W-specific insertions are shown on ChrW in zoom. The deletions and insertions are mainly transposable elements.

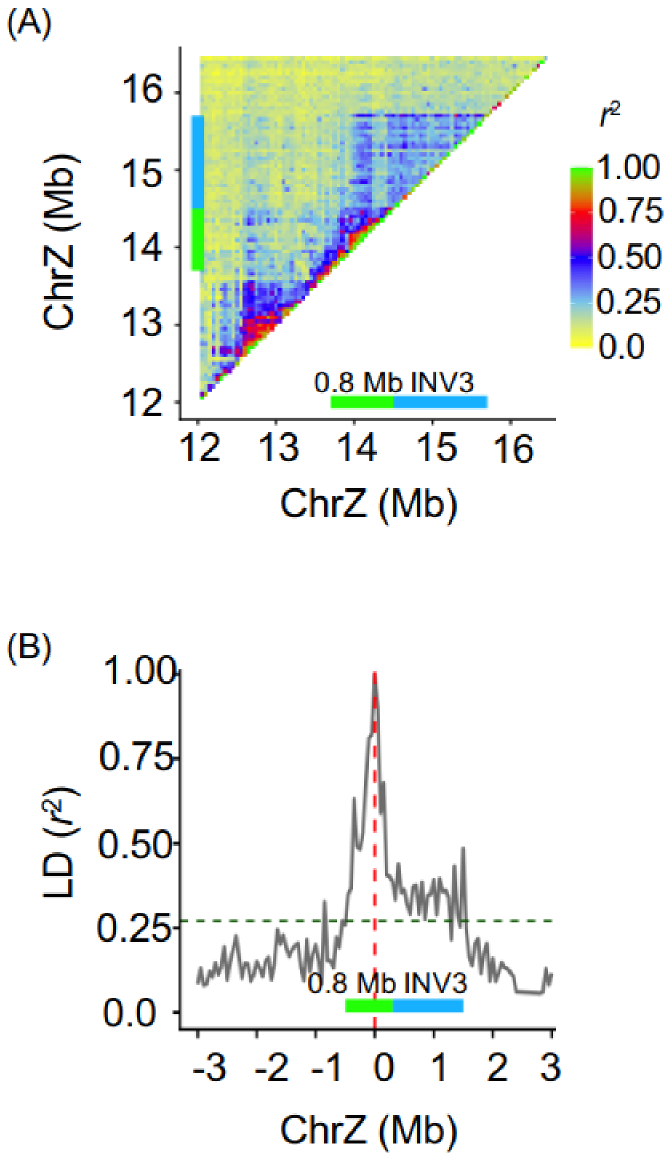


**Supplementary Fig. 14. (A)** Linkage disequilibrium (LD) around the 0.8 Mb and INV3 regions in the reference genome of only ZZ males of *Scomber japonicus*. LD heatmaps and maximum $r^2$ values are shown. **(B)** LD decay plot with a 100-kb window from the focal position of 14.2 Mb on ChrZ.

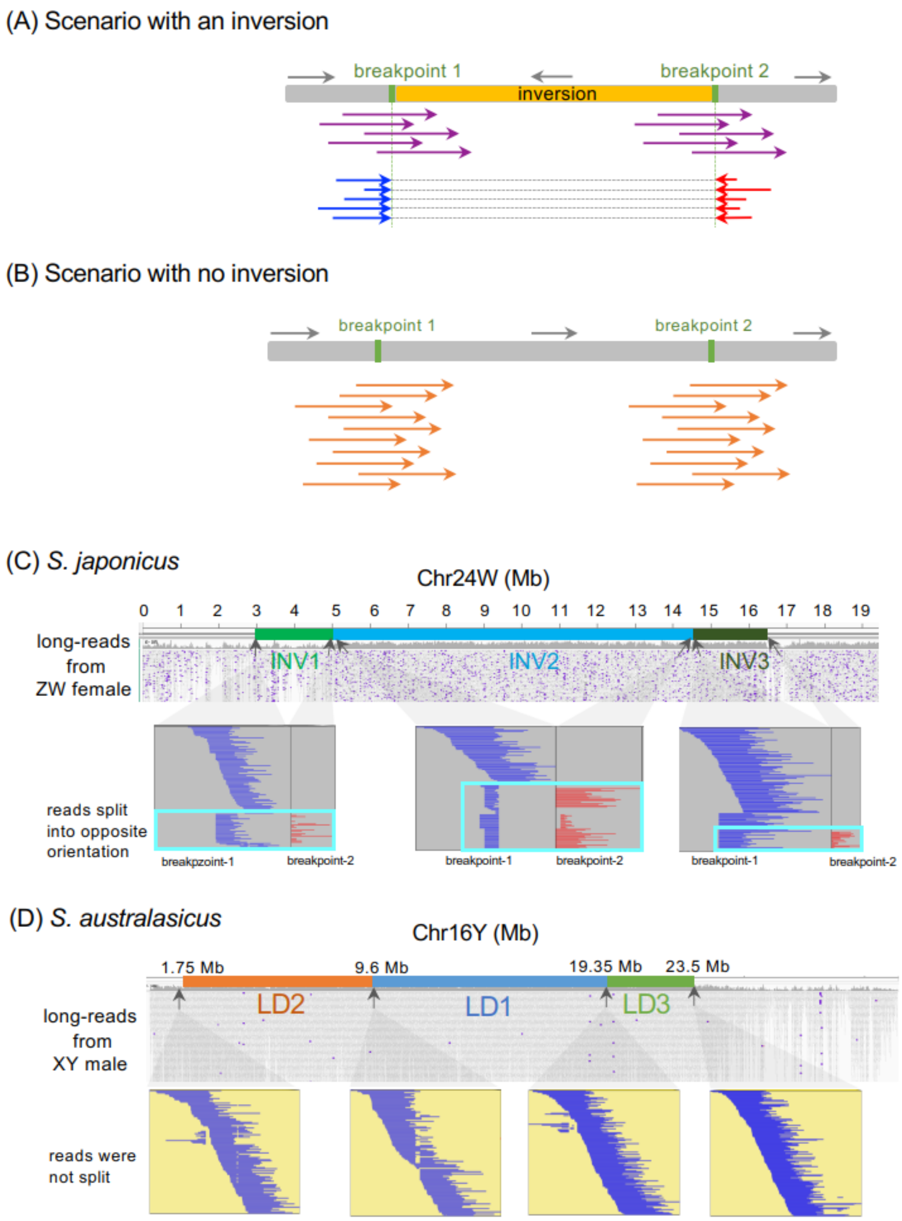


**Supplementary Fig. 15.** Inversions detected in the split read analysis. **(A)** Illustration of a scenario with inversion. When long reads from heterogametic individuals are mapped onto a chromosome with an inversion, reads with a noninverted orientation are split at the two borders of the inversion, and the read segments are mapped in the opposite orientation (blue and red arrows), whereas reads with an inverted orientation (purple arrows) are not. **(B)** Illustration of a scenario without inversion. No split reads are observed (orange arrows). **(C)** Alignment results of the HiFi long reads of the ZW individual onto Chr24W. Top panel: IGV image of the aligned reads. The breakpoints are indicated as grey arrows. Bottom panel: Reads from Z chromosomes (in cyan rectangles) were split into two parts at the breakpoints for each of the inversions. **(D)** Alignment results of the HiFi long reads of the XY individual onto Chr16Y. Top panel: IGV image of the aligned reads. The boundary of each high-LD region is indicated with grey arrows.

Bottom panel: Split reads were not observed near the boundaries for either of the high-LD regions.

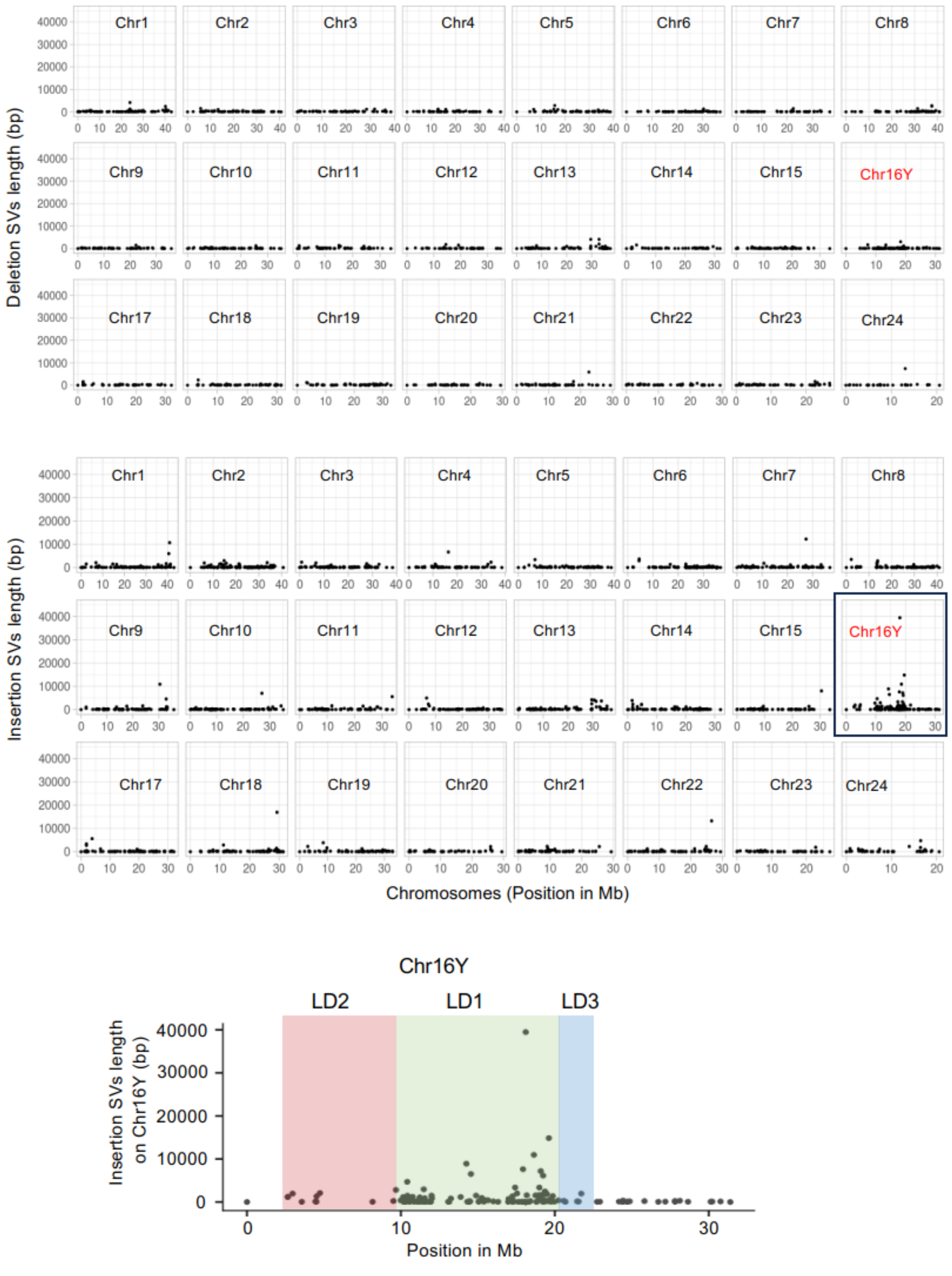


**Supplementary Fig. 16.** Insertion and deletion structural variants (SVs) are enriched on Chr16Y. Structural variant calling using haplotype-resolved genome assemblies and read-based alignments. For read-based structural variant calling, we used the HiFi log-reads from two male individuals and one female individual and the Oxford Nanopore long-reads from one male and one female individual. We mapped those reads onto the SAM-hap1 reference genome. We identified the male-biased deletion and insertion SVs (> 50 bp) that are heterozygous for males but homozygous for females. **(A)** The identified deletion SVs are shown for 24 chromosomes in the top plot, and the insertion SVs are shown in the bottom plot. The lengths of the SVs are shown on the y-axis, and the chromosomes are shown on the x-axis. **(B)** Y-specific insertions are shown on Chr16Y in zoom. The deletions and insertions are mainly transposable elements.

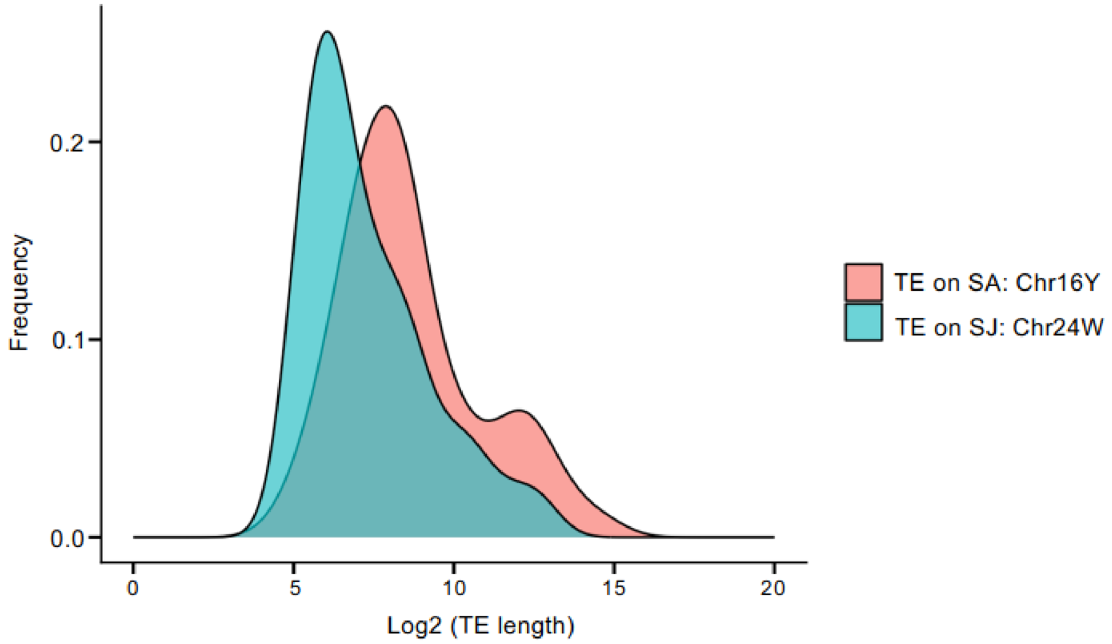


**Supplementary Fig. 17.** Distribution of the TE length on the Chr16Y- and Ch24W-specific insertions identified in *S. japonicus* (SJ) and *S. australasicus* (SA), respectively.

### Supplementary methods

#### *De novo* reference genome assemblies of a ZW *Scomber japonicus* and a XY *S. australasicus*

Draft genomes were initially assembled with HiFi reads by hifi-asm (v. 0.16.1)[5,6] integrated with Omni-C data with options of s=0.40, D=10, and N=200. The Omni-C data were further utilized for scaffolding. Reads were mapped onto the haplotype-resolved contigs produced by hifi-asm using Juicer (v. 1.6)[7] with options of s=none. These contigs were scaffolded using read pairs uniquely mapped onto either of the contig pairs to generate a haplotype-resolved reference genome using 3DDNA (v. 180922)[8] with the options r=0 and editor-repeat coverage=15. Visual inspection of the Omni-C contact maps and manual correction of the misassemblies and misscaffoldings (e.g., inversions and translocations) were performed using Juicebox (v. 1.11.08)[9]. Moreover, scaffolds that could not be assigned to any of the 24 major chromosomes were manually joined into large superscaffolds based on the chromatin contact frequency. The completeness of each haploid genome assembly was assessed using gVolante (v. 1.2.1)[10] with BUSCO (v. 3.0.2)[11] and the orthologue gene set of Actinopterygii.

#### Chromosome-level genome assemblies for additional *S. japonicus* and *S. australasicus* individuals

Phased contigs were generated using hifi-asm with the default settings. The phased contigs were then scaffolded at the chromosome scale using Ragtag[12], guided by the reference genomes of the ZW *S. japonicus* and the XY *S. australasicus* individuals constructed as described above.

#### Reassembling *S. japonicus* and *S. scombrus* genomes using public datasets

Haplotype-resolved contigs of these fish were assembled using long-read HiFi and Arima Hi-C sequences using hifi-asm (v. 0.16.1) with options of s=0.40, D=10 and N=150. These haplotigs were further scaffolded using the Arima Hi-C data. First, sequence reads of Arima Hi-C were mapped onto the contigs using Juicer (v. 1.6) with the option s=Arima. Unmapped or low-quality reads, as well as reads derived from PCR duplicates, were filtered out as described above. The produced BAM file was manually split into two segments, each containing reads mapped onto the primary and alternative assemblies (haplotypes), and then sorted using SAMtools (v. 1.21). These contigs were scaffolded by subjecting these haplotype-specific BAM files to YaHS (v. 1.2)[13] with the option –no-contig-ec, and Hi-C contact maps were generated using Juicer's pre command. Misassembled and misscaffolded sequences were manually corrected using Juicebox (v. 1.11.08).

#### Gene and functional annotations

Gene and functional annotations of the haploid-resolved *S. japonicus* (SJF-hap1) and *S. australasicus* (SAM-hap1) assemblies were performed using a combination approach of RNA-seq-based and *ab initio* prediction with the Augustus/BRAKER pipeline (v. 3.0.3)[14]. Transcriptomes were collected from six tissues, including the eye, liver, muscle, kidney, spleen, and brain, obtained from one individual per sex per species. Total RNA was extracted with the conventional spin-column method, and sequencing libraries were constructed using a TruSeq stranded mRNA Library Preparation Kit (Illumina). These libraries were sequenced on a NovaSeq 6000 instrument with 150PE. RNA extraction, library construction, and sequencing were performed by Macrogen Japan. Sample information and raw reads are summarized in Supplementary Table S15. These RNA-seq reads were mapped onto the respective reference genomes (the ChrW-containing haploid genome for *S. japonicus* and the ChrY-containing the haploid genome for *S. australasicus*) using HISAT2 (v. 2.1.0)[15] in the setting of --wrapper basic-0 -k 1 --no-discordant --no-mixed -p 20 –dta and then sorted based on the genomic position using SAMtools (v.1.21). The mapping results were used for evidence-based gene annotation in conjunction with the annotated protein sequences from fScoJap1 (GCF_027409825.1)[2].

#### Repeat annotation

A *de novo* transposable element database was constructed for each species using RepeatModeler (v. 2.0.2)[16] with RECON[17], RepeatScout[17,18] and CD-HIT-EST[19]. Full-length, long-terminal repeat retrotransposons were identified using LTRharvest and LTRdigest[20,21]. These databases were combined with the nonredundant de novo TE library of *Scombrini* available from Dfam (v. 3.8) (https://www.dfam.org/releases/Dfam_3.8/families/FamDB/). Interspersed and simple repeats were annotated for each genome using RepeatMasker (v. 4.1.6) (http://www.repeatmasker.org) with the following parameters: -pa 12 -s -no_is -u -noisy -xm -a -xsmall -uncurated

#### Inference of telomere and centromere positions

Telomeric repeats (TTAGGG) were scanned using TeloExplorer implemented in quarTeT (v. 0.2) (https://github.com/aaranyue/quarTeT) with the -c animal option. The putative centromere positions were also identified through the following steps. First, large arrays of tandem repeats were visualized on the self-vs.-self-aligned dot plot drawn for each chromosome using self-BLAST results with the option word_size 100; centromere regions are assumed to be characterized by a large array of tandem repeats dense on the diagonal. Centromeric repeats were subsequently retrieved using quarTeT/CentroMiner with -TE, -gene, and -m 400 options. Finally, CpG methylation landscapes were inferred from PacBio HiFi kinetic data[22]; signatures of DNA hypomethylation emerged in the centromeric regions because of the formation of kinetochore complexes. Consensus read construction was performed using pbccs (v. 6.4) (https://github.com/PacificBiosciences/ccs) with the parameter --hifi-kinetics, and then, the probabilities of CpG methylation were estimated per base using jasmine (v. 2.0.0) (https://github.com/pacificbiosciences/jasmine). HiFi reads annotated with CpG methylation probabilities were mapped onto the primary haploid genome using pbmm2 (v. 1.13.1) (https://github.com/PacificBiosciences/pbmm2) with the default setting. The per-base CpG methylation probability was estimated from the resulting BAM file by employing aligned_bam_to_cpg_scores (v. 2.3.0) (https://github.com/PacificBiosciences/pb-CpG-tools) with the default setting. The combination of these three approaches, i.e., self-aligned dot plots, centromere repeat mining, and CpG methylation landscapes, allowed us to identify putative active centromeric satellite arrays >100 kb in size.

Conserved centromeric satellite DNA shared between the two pacific *Scomber* species was identified from tandem repeats (TRs) detected using quarTeT/CentroMiner. A total of 830 and 803 TRs were identified in *S. japonicus* (SJF-hap1) and *S. australasicus* (SSM-hap2), respectively, with repeat unit lengths ranging from 99401 bp and 98392 bp, respectively. Clustering with CD-HIT at 90% sequence similarity yielded 250 clusters for *S. japonicus* and 243 clusters for *S. australasicus*. The repeat copy number for each cluster was quantified by BLAST searches against the respective genome assemblies. The 100 most abundant TR clusters from each species were further clustered at 70% sequence similarity, resulting in 15 clusters in *S. japonicus* and 21 in *S. australasicus*. These 36 clusters were then classified into families based on 80% sequence similarity using reciprocal BLASTN alignment. This analysis revealed seven families of conserved centromeric satellite DNA shared by both species (Supplementary Table S3).

#### Construction of a phylogenetic tree of the genus *Scomber* using single-copy orthologues

Species phylogeny within the genus *Scomber* was inferred using single-copy orthologous gene sequences. *S. colias* was included in this analysis, and two species from the sister genus *Thunnus*, i.e., *T. maccoyii* and *T. thynnus*, were used as outgroups. Protein sequences of *S. japonicus*, *S. australasicus*, and *S.*

*scombrus* were obtained as described above, while those of *S. colias* (GCA_021039105.1; GCF_963691925.1), *T. maccoyii* (GCF_910596095.1), and *T. thynnus* (GCF_963924715.1) were retrieved from NCBI databases.

A total of 6,400 single-copy orthologous sequences were identified using OrthoFinder (v.2.4)[23] with the default settings, including option -M. Alignments of each orthologue pair were obtained using MUSCLE (v. 3.8.31)[24] with the default settings. Alignment gaps were trimmed using TrimAl (v. 1.2)[25] with a threshold of 0.85 (-gt 0.85), and all sequences were concatenated per species using joint_alignment.py implemented in supermatrix script (https://github.com/wrf/supermatrix). The supermatrix fasta file and partition file are available in figshare (https://doi.org/10.6084/m9.figshare.31977930). The phylogenetic relationships among species were inferred using IQ-Tree (v. 1.6.12)[26] with the parameters -bb 10000, -nt AUTO, and -st AA. The best-fit molecular evolutionary model (Q. bird+F+I+R7) was selected by ModelFinder to construct a phylogenetic tree within the IQ-Tree.

#### Analysis of synteny among haploid genomes

Large-scale interspecies chromosomal rearrangements were assessed through an analysis of synteny among haploid genomes between and within *S. japonicus*, *S. australasicus*, and *S. scombrus*. The chromosome-scale reference genomes of two outgroup species, *T. maccoyii* (GCF_910596095.1) and *T. thynnus* (GCF_963924715.1), were included. Each genome was reciprocally mapped using Minimap2 (v. 2.21)[27] with the options -ax asm5 for within-species alignments and -ax asm10 for between-species alignments[28]. Larger structural rearrangements such as inversions, duplications and translocations were identified using SyRI (v. 1.6.3)[29] and visualized using plotsr (v. 1.1.0)[30]. The orientation of these variations was inferred from sequence collinearity between the two *Thunnus* species. Large inversions (>1 Mb) were also visualized as dot plots generated using BLASTN with a word_size of 30. We also utilized dot plots to identify the repeat arrays at the breakpoints.

#### Structural variant detection using long-read sequences

Read discontinuities were analysed using PacBio HiFi long reads generated for genome assemblies and Oxford Nanopore long reads generated using the PromethION platform (Nanopore Technology, UK), including a total of 3 females and 2 males for *S. japonicus* and 2 females and 3 males for *S. australasicus* (Supplementary Table S14).

Genomic DNA for Nanopore sequencing was obtained from whole-blood samples of wild-caught individuals—one sample for each sex of each species: a ZW female and a ZZ male (*S. japonicus*) commercially caught in the offshore area of Oita, Japan, and an XX female and an XY male (S. *australasicus*) commercially caught in the offshore area of Nagasaki, Japan. Genomic DNA extraction and sex identification were performed as described above. DNA fragments smaller than 10 kb were depleted using the Short Read Eliminator XS kit (Circulomics, Baltimore, MD, USA) to increase the read length. A 1D Ligation Sequencing Kit (LSK109) was used for library construction, and each individual was sequenced on a single flow cell (FLO-PRO002). Library preparation and sequencing were performed by GeneBay Inc. These long reads were mapped onto either SJF-hap1 or SAM-hap1 using minimap2 with option -ax map-hifi for the HiFi reads and -ax map-ont for the Nanopore reads.

Reads with a mapping quality of less than 30 and secondary alignment were removed, and the resulting BAM files were sorted using SAMtools. BAM files were visualized to identify patterns of read discontinuity near inversion breakpoints. Breakpoint coordinates were further refined by inspecting the BAM files for reads split across inversion junctions, where one part of the read aligned to one breakpoint and the other aligned to the opposite breakpoint in the reverse orientation. We characterized these events by focusing on 100-kb genomic regions flanking the proximal and distal breakpoints of each inversion.

We annotated long-read alignments using Ribbon (v. 2.0)[31] to examine split reads supporting inversion structures.

We used SVIM (v. 2.0.0) (https://github.com/eldariont/svim)[32] with the option alignment to call SVs. We then merged the species-specific VCF files using SURVIVOR (v. 1.0.7) (https://github.com/fritzsedlazeck/SURVIVOR)[33] with the following parameters: 1000, 1, 1, 0, 0, and 50. These parameters include the following: a maximum allowed distance between SV breakpoints for merging of 1,000 base pairs; the need for at least one supporting caller to consider the SV; the SV type; the neglect of the SV strand orientation; the inability to estimate the breakpoint distance based on the SV size; and a minimum SV size of 50 bp. Structural variant calling within highly repetitive regions is highly challenging; thus, SVs primarily composed of simple repeats were excluded from this analysis. Sex-specific SVs (> 50 bp) were identified using a Perl script, Find_sex_loci_from_GT.pl (https://github.com/lyl8086/find_sex_loci)[34] with option -c 3; SVs heterozygous (0/1) in females and alternative homozygous (1/1) in males were extracted for *S. japonicus* with a female heterogametic system, whereas those heterozygous (0/1) in males and alternative homozygous (1/1) in females for *S. australasicus* were extracted with a male heterogametic system.

#### Linkage disequilibrium

We also computed the correlations ($r^2$) in the genotypes between each SNP pair across the sex chromosomes for males and females separately to understand the occurrence of recombination suppression between gametologous sex chromosomes. We used the geno-r2 function of VCFtools (v. 1.16)[35]. The mean and maximum $r^2$ values within a 50-kb window were calculated using the emerald2windowldcounts.pl script (https://github.com/owensgl/haploblocks).

#### Assessment of intraspecific structural variations through a local principal component analysis (PCA)

A local PCA was performed using low-coverage whole-genome sequence (lcWGS) data obtained from wild-caught individuals to examine intraspecific heterogeneity in genomic structure, including structural variations such as chromosomal inversions, in the two Pacific *Scomber* species. A total of 281 (136 males and 145 females) individuals of *S. japonicus* captured from Tatayama, Fukui, and Oita (Japan) and a total of 188 (96 males and 92 females) individuals of *S. australasicus* captured from Tateyama and Nagasaki (Japan) were used (Supplementary Table S5). The phenotypic sex of each individual was determined visually. DNA was extracted from the caudal fin using a Gentra Puregene tissue kit (Qiagen) according to the manufacturer's instructions. The sequencing library was constructed using a Twist 96-plex library preparation kit (Twist Bioscience, USA) according to the manufacturer's protocol, and sequencing was performed on a single lane of a NovaSeq 6000 S4 flow cell (PE 150) at Rhelixa, Inc. (Tokyo, Japan), except for Oita samples, which were sequenced on a single lane of DNBSEQ G400-RS at GeneBay, Inc.

Genotypes of these individuals were inferred from posterior genotype probabilities using individual allele frequencies as prior information using PCAngsd (v. 0.99)[36], as described previously[37]. Quality filtering was applied to the sequence data using Atria (v. 3.1.1)[38] with the default parameters, followed by demultiplexing using the DemuxFastqs function of fgbio (v. 2.0.1) (https://github.com/fulcrumgenomics/fgbio), with the read structures and sample index specified as --read-structures 8B12S+T 8S+T --metrics index.file, where the index.file included sample index sequences retrieved from the manual of the library construction kit. Filtered reads were mapped onto species-specific reference genomes (SJF-hap1 for *S. japonicus* and SAM-hap1 for *S. australasicus*) using BWA-MEM[39] with the default settings. The generated SAM files were converted to BAM format and sorted based on the genomic position using SAMtools. The sorted BAM files were subjected to local realignments using GATK (v. 3.8) IndelRealigner[40]. The input beagle format file was generated by ANGSD (v. 0.94)[41] in GATK genotyping mode (-GL 2) with the following options: -remove_bads 1, -

trim 0, -minMapQ 20 and -minQ 20 to remove low-quality reads and bases; -skipTriallelic to retain only biallelic SNPs; -uniqueOnly 1 to exclude reads not uniquely mapped onto the reference; -only_proper_pairs 1 to keep only properly mapped pairs; -minMaf 0.05 to exclude variants whose minor allele frequency was less than 0.05; -minInd 100 to retain variant sites where at least 1 read was mapped onto the position for more than 100 individuals; and -doGlf 2 and -doMajorMinor to generate beagle format genotype likelihoods files. Genotype calling was performed on the output beagle file using PCAngsd with the options -sites_save, -post_save, and -e 3 while principal component analysis (PCA) was conducted; the number of eigenvalues (-e) was set to 3 for *S. japonicus* and -e 2 for *S. australasicus* according to the number of populations. The output file was converted to a VCF format file using beagle2vcf.awk (https://github.com/ariloytynoja/saimaa-population-structure/blob/main/beagle2vcf.awk).

SNP filtering was subsequently performed for each species using VCFtools (v. 1.16). SNP sites that met the following criteria were retained: a minor allele frequency (MAF) greater than 0.01 (--maf 0.01), biallelic (--min-alleles 2 --max-alleles 2), and minimum genotyping rates of 0.8 for *S. japonicus* (--max-missing 0.8) and 0.6 for *S. australasicus* (--max-missing 0.6). A total of 14,086,035 and 3,857,157 SNPs were obtained from *S. japonicus* and *S. australasicus*, respectively. The resulting VCF files were converted to BCF format using the convert command of BCFtools (v. 1.10.2)[42].

A local PCA was performed using the lostruct R package (https://github.com/petrelharp/local_pca)[43]. Local PCA coordinates were computed using the eigen_windows function for each nonoverlapping 10-kb window. Pairwise distances between PCA maps were calculated based on the top two principal components using the pc_dist function with the default parameters. The resulting distance matrix was subjected to multidimensional scaling (MDS) with the cmdscale function to visualize genomic regions with structural variation. Genomic loci exhibiting structural variants, such as inversions, appear as outliers when MDS values are plotted against chromosomal positions. Finally, locus-specific PCA was performed using the R package SNPRelate (v. 1.40)[44,45].

Localized heterogeneity is often associated with polymorphic chromosomal inversions. In such cases, individuals typically cluster into two or three genotype groups: (i) homozygotes for the standard haplotype, (ii) heterozygotes carrying both standard and inverted haplotypes, and (iii) homozygotes for the inverted haplotype. A cluster consisting of heterozygotes is supposed to display higher average heterozygosity than a cluster consisting of homozygote groups. In addition, the regions harbouring polymorphic inversions form linkage disequilibrium (LD) blocks. We therefore determined the MDS outlier regions as intraspecific inversions if the region exhibited an LD block; individuals were separated into two or three distinct clusters within the region, and one cluster showed increased heterozygosity if the genotype frequency followed the Hardy–Weinberg equilibrium ($p > 0.05$). Individual clustering was performed using *k*-means clustering implemented in R as the kmean function, with the k value selected using the elbow method. The average heterozygosity across the outlier region was calculated per individual using PLINK (v. 2)[46]. Pairwise genotype correlations ($r^2$) between SNPs were computed across chromosomes using the geno-r2 function in VCFtools (v. 1.16). Conformity to HWE was tested for each region with the HWE.test function in the R package genetics (v. 1.3.8.1.3)[47]

#### Sex chromosome scanning for *S. japonicus* and *S. australasicus*

Sex-associated SNP markers for *S. japonicus* (Jpn1 and Jpn2) and *S. australasicus* (Aus1 and Aus2) have been identified with a reference-free approach using pooled sequencing data[48]. The genomic positions of these sex diagnostic markers were located using a BLAST search against the haplotype-resolved reference genomes of each species to identify the sex chromosomes of *S. japonicus* and *S. australasicus*. The aforementioned VCF file generated from low-coverage whole-genome resequencing data was subjected to a genome-wide association study (GWAS) through Fisher's exact test under a dominant model using PLINK (v. 1.07) to further determine the entire SD region in the sex chromosome[49]. The genome-wide significance threshold was determined using a Bonferroni correction based on the total number of SNPs tested: $P < 0.05/N$, where $N$ is the total number of SNPs tested. Manhattan plots were constructed using the R package qqman[50]. Sex-specific SNPs within the SD region were identified using the Perl script find_sex_loci_from_GT.pl (https://github.com/lyl8086/find_sex_loci). The differences in genomic diversity between the sexes were scanned using Weir and Cockherham's $F_{ST}$[51] within a 10-kb nonoverlapping window using VCFtools.

#### Localization of the sex-determining region in *S. scombrus*

RAD-seq reads were filtered using Trimmomatic (v. 0.36) and mapped onto the primary haploid genome of *S. scombrus* (SSM-hap1) using BWA-MEM with the default settings. Genotype calling was performed using GATK HaplotypeCaller (v. 3.8)[40] with the default settings. The obtained SNPs were analysed using VCFtools (v. 0.1.14) with the settings, --max-missing 0.3 and --maf 0.1, keeping only biallelic SNPs with a minimum depth of 5. These SNPs were subjected to GWAS, SNP density plotting, and site $F_{ST}$ using the aforementioned protocols; however, we could not detect clear peak pattern from these analyses. Thus, we adopted a reference-free approach using RADSex (v. 1.1.0) (https://github.com/SexGenomicsToolkit/radsex)[52]. Marker tables, including the depth of coverage, were generated from filtered RAD-seq reads using the *process* command. The marker distribution across sexes was calculated with the *distrib* command, and markers significantly associated with phenotypic sex were identified using the *signif* command with the option d=5. A single marker was detected almost exclusively in males and was significantly associated with sex (adjusted $P < 0.05$, $\chi^2$ test with the Bonferroni correction). We performed a BLASTn search using the sequence of the sex-associated marker as the query against the haploid assemblies of *S. scombrus* to identify the male heterogametic region in the haploid reference genome and confirmed that SSM-hap1 contains the male heterogametic region. We further mapped all the RADSex markers with a map command (implemented in RADSex) on SSM-hap1 and constructed a Manhattan plot.

The locations of heterogeneous SD regions were scrutinized by performing a depth-of-coverage analysis using whole-genome resequencing data. We obtained resequencing data from ten individuals of *S. scombrus* from the NCBI SRA database (PRJNA253681). Reads were trimmed with Trimmomatic (v. 0.36) using the aforementioned protocol. These filtered reads were mapped onto the SSM-hap1 assembled genome of *S. scombrus* using BWA-MEM. The resulting BAM files were sorted, and PCR duplicates, reads with low mapping quality ($q < 10$), and ambiguously mapped reads were excluded using SAMtools (v. 1.15.1). The depth of coverage was determined with Mosdepth (v. 0.3.0)[53] per 1-kb window and normalized to the genome coverage. The genotypic sex of these individuals was determined by the presence (male) or absence (female) of reads mapped onto the SDR.

#### Genetic diversity analysis and neutrality test

Genetic diversity between two sample groups was assessed using Weir and Cockherham's $F_{ST}$[51] within a 100-kb nonoverlapping window. Neutrality tests (Tajima's D)[54] were performed separately for each sample group within a 100-kb nonoverlapping window. VCFtools was used for these analyses.
